# Selective epigenetic regulation of IFN-γ signature genes by JAK inhibitor in inflammatory diseases

**DOI:** 10.1101/2024.08.05.606293

**Authors:** Geunho Kwon, Yebin Park, Keunsoo Kang, Kyuho Kang

**Affiliations:** Department of Biological Sciences and Biotechnology, Chungbuk National University, Cheongju 28644, Republic of Korea; Department of Microbiology, College of Science & Technology, Dankook University, Cheonan 31116, Republic of Korea

## Abstract

This study addresses the molecular mechanisms behind JAK inhibitors’ (JAKi) epigenetic regulation of IFN-γ-induced genes in macrophages, key players in controlling inflammation in chronic diseases such as rheumatoid arthritis (RA) and COVID-19. Utilizing transcriptomic and epigenomic approaches, we reveal that JAKi selectively regulate gene programs in IFN-γ-activated human macrophages. Our single-cell analysis identified high expression of IFN-γ signature genes in macrophages from RA and severe COVID-19 patients. JAKi suppressed some inflammatory genes, while a subset remained unresponsive to JAK-STAT inhibition. We discovered that JAKi selectively target IRF-STAT-dependent open chromatin regions, leaving AP-1-C/EBP-regulated genes with open chromatin unaffected. Some JAKi-insensitive genes could be inhibited by JNK inhibitors in IFN-γ-primed macrophages. Our analysis also identified JAKi-sensitive and - insensitive IFN-γ signature genes in RA patients resistant to MTX treatments and COVID-19 vaccinated donors, highlighting JAKi’s therapeutic potential and risks. These findings uncover new JAKi responsiveness mechanisms through epigenomic changes in IFN-γ-primed macrophages, advancing our understanding of inflammation regulation in chronic diseases.

**Teaser:** Discover how JAK inhibitors selectively regulate gene programs, shedding light on inflammation control in chronic diseases.

## Introduction

Inflammation is a multifaceted immune response triggered by various stimuli such as pathogens and damaged cells. Uncontrolled inflammation can lead to the development of several inflammatory diseases including lupus, rheumatoid arthritis (RA), inflammatory bowel disease (IBD), and COVID-19 (*1–3*). These conditions are marked by abnormal immune responses and cytokine overproduction, particularly TNF and IL-6 (*4–6*). Macrophages are central to this response, exhibiting dynamic changes in their transcriptomic, epigenomic, and metabolic programs in response to environmental signals (*7–9*). These cells display heterogeneous phenotypes shaped by signals from the tissue microenvironment (*10–15*). The presence of an “interferon signature” within innate immune cells, including macrophages, is a hallmark of inflammatory diseases (*16–18*). Pathogenic macrophages in inflammatory diseases are activated by the combined effects of IFN-γ and other cytokines (*18, 19*). Gaining deeper insights into the molecular mechanisms associated with interferon signatures is crucial to understand the underlying mechanisms of inflammatory diseases.

Interferon-γ (IFN-γ) is a pro-inflammatory cytokine that elicits activation of macrophages and drives a pro-inflammatory “M1-like” phenotype (*16*). IFN-γ priming causes epigenetic modifications and reprogramming of metabolic pathways in macrophages, resulting in enhanced expression of “M1-like” inflammatory genes and reduced expression of “M2-like” anti-inflammatory genes (*20, 21*). This “epigenomic signature” of IFN-γ-primed macrophages is linked to the onset of inflammatory diseases (*22, 23*). Activation of the JAK-STAT signaling pathway in IFN-γ-primed macrophages leads to the phosphorylation of both STAT1 and STAT3 (*24–27*). The competition between STAT1 and STAT3 for binding to specific *cis*-regulatory elements can influence gene expression and the inflammatory response. A deeper understanding of the interactions between these signaling pathways and their epigenetic regulation by STATs and other transcription factors in the context of inflammatory diseases is crucial for the development of effective therapies.

Inflammatory diseases, such as RA and COVID-19, are commonly treated with JAK inhibitors (JAKi), drugs that target the JAK-STAT signaling pathway (*28, 29*). Repeated treatment with JAKi has been shown to repress IFN-signature genes and is associated with the modification of chromatin accessibility at *cis*-regulatory elements (*30*). The presence of shared transcriptomic profiles, including interferon signatures, among various inflammatory diseases suggests the possibility of a common treatment approach (*18*). Tofacitinib, an oral JAKi, effectively blocks the activity of JAK1 and JAK3, leading to a reduction in pro-inflammatory cytokine production and a balanced immune response (*31*). In some cases, tofacitinib is used in combination with other disease-modifying anti-rheumatic drugs (DMARDs), or as a last resort for patients who do not respond to other treatments such as methotrexate (MTX) and TNF inhibitors (*32*). The recent research highlights the significance of identifying different clinical response profiles to optimize pharmacotherapy (*33*). The ability of JAKis to regulate the immune response, specifically the cytokine storm seen in severe COVID-19 cases, suggests their potential as a treatment option for COVID-19 (*34, 35*). However, safety concerns regarding JAKis, particularly with regards to cardiovascular and cancer risk, have been raised in older RA patients who have cardiovascular risk factors (*36, 37*). To fully leverage the promise of JAKis in treating autoimmune and inflammatory diseases like RA and COVID-19, a deeper understanding of the molecular mechanisms underlying their regulation of the inflammatory response is necessary.

In this study, we aimed to uncover the molecular mechanisms by which JAKi, drugs widely used to treat inflammatory diseases like RA and COVID-19, regulate the expression of IFN-γ-induced genes in human macrophages. We combined transcriptomic and epigenomic methods to explore how JAKi modulate gene expression through epigenetic changes in IFN-γ-primed human macrophages. Our analysis showed differential patterns of IFN-γ signature genes based on JAKi sensitivity. JAKi-sensitive genes displayed reduced chromatin accessibility at cis-regulatory regions enriched for IRF-STAT motifs, while JAKi-insensitive genes maintained open chromatin regions with heightened AP-1 and C/EBP motifs. We discovered that JNK inhibition suppresses JAKi-insensitive genes in IFN-γ-stimulated human macrophages. Our transcriptomic analysis of RA patients resistant to MTX and COVID-19 vaccinated donors revealed the presence of both JAKi-sensitive and JAKi-insensitive IFN-γ signature genes in macrophages, emphasizing the potential therapeutic benefits and adverse effects of JAKi treatment. This study provides new insights into the previously unexplored mechanisms of JAKi responsiveness through epigenomic changes in IFN-γ-primed human macrophages and deepens our understanding of the regulatory landscape in inflammatory diseases.

## Results

### Transcriptomic signatures of macrophages in inflammatory diseases

To characterize the IFN-γ signature genes in human macrophages, we conducted a transcriptomic analysis and compared the gene expression profiles of IFN-γ-primed macrophages with those stimulated by TNF and LPS (*21, 38, 39*). Our analysis revealed 235 genes (11.5%) that were specifically upregulated by IFN-γ, primarily related to metabolic processes such as cholesterol biosynthesis, ATP metabolism, and proteasome activity (Fig. 1, A and B). Conversely, 81 genes (6.1%) were specifically downregulated by IFN-γ, with functions involved in extracellular matrix organization and degradation (fig. S1, A and B). Moreover, 181 genes (8.9%) were commonly upregulated by all three stimuli, and these genes were linked to the functions of inflammatory response (Fig. 1, A and B). Additionally, 106 genes (8%) that were commonly downregulated by the three stimuli played a role in iron ion transport and receptor-mediated endocytosis (fig. S1, A and B).

**Fig. 1.**
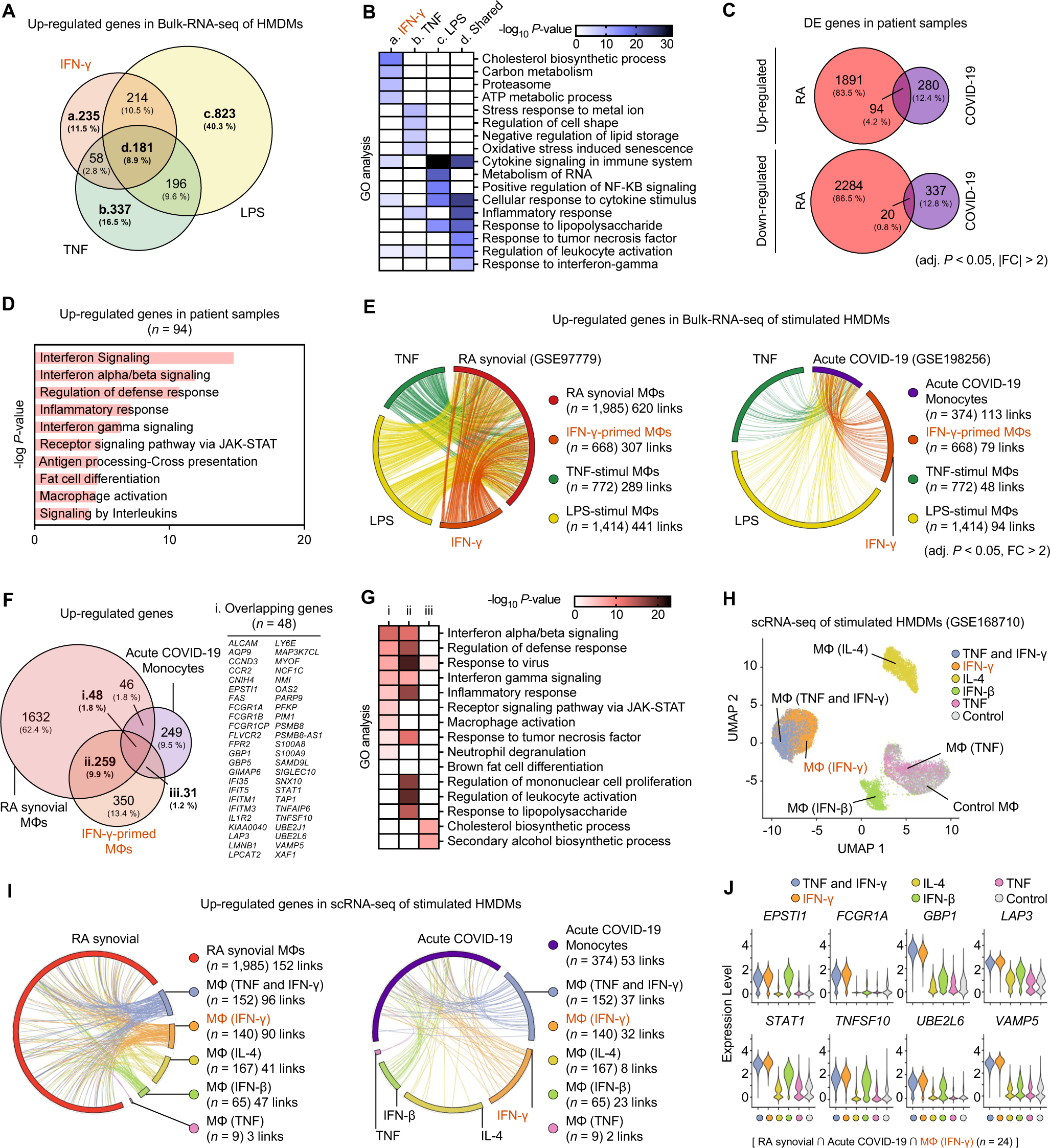
IFN-γ signatures in inflammatory macrophages unveiled through transcriptome profiling. **(A)** Venn diagrams represent upregulated genes in human monocyte-derived macrophages (HMDMs) upon stimulation with IFN-γ, TNF, or LPS, as identified from RNA-seq data of three independent studies (GSE98368, GSE120944, and GSE100382). EdgeR was used for analysis (FDR adjusted P < 0.05, fold change > 2), and transcripts per million (TPM) values were filtered to be greater than 4. **(B)** Gene ontology (GO) analysis of upregulated genes specifically or commonly induced by each stimulus. **(C)** Venn diagrams illustrate differentially expressed (DE) genes in rheumatoid arthritis (RA) synovial macrophages and acute COVID-19 monocytes. **(D)** GO analysis of overlapping DE genes in RA synovial macrophages and acute COVID-19 monocytes. **(E)** Circos plots show upregulated genes linked between RA synovial macrophages or acute COVID-19 monocytes and HMDMs primed or stimulated with cytokines, color-coded based on priming or stimulation. **(F)** Venn diagram of genes upregulated in RA synovial macrophages, acute COVID-19 monocytes, and IFN-γ-primed HMDMs, with shared genes across all data sets indicated on the right. **(G)** GO analysis of upregulated genes common to RA synovial macrophages, acute COVID-19 monocytes, and IFN-γ-primed HMDMs, using the “i”, “ii”, and “iii” genes identified in (F). **(H)** Single-cell RNA-seq (scRNA-seq) data analysis of stimulated and unstimulated HMDMs, color-coded and labeled based on cytokine stimuli. **(I)** Circos plots display overlapping DE genes among RA synovial macrophages, acute COVID-19 monocytes, and cytokine-stimulated HMDMs, with linked genes color-coded according to stimulation conditions in scRNA-seq. **(J)** Violin plots depict gene expression in stimulated and unstimulated HMDMs, using positively correlated genes identified in each data set. Metascape was utilized for GO analysis.

We investigated the correlation between IFN-γ signature genes and inflammatory diseases by analyzing transcriptomic data from RA synovial macrophages (GSE97779 (*21*)) and monocytes from COVID-19 patients (GSE198256 (*40*)) (fig. S1, C-E). Our analysis revealed common enhancement of 94 genes related to IFN-γ-related biological functions and regulation of the interferon signaling and JAK-STAT signaling pathways in both RA synovial macrophages and COVID-19 monocytes (Fig. 1, C and D). Notably, RA synovial macrophages showed the strongest positive correlation with IFN-γ-primed macrophages, with 46.0% of genes upregulated and 52.6% downregulated in response to IFN-γ stimulation (Fig. 1E and fig. S1F). Moreover, we found that among the three stimuli (IFN-γ, TNF, and LPS), the genes upregulated by IFN-γ displayed the strongest correlation with genes upregulated in COVID-19 monocytes, with 11.8% of these genes showing such correlation, compared to TNF (6.3%) and LPS (6.6%) (Fig. 1E). In contrast, the correlation between IFN-γ-downregulated genes and COVID-19 monocytes was less prominent (fig. S1F). To further examine the transcriptional landscape of inflammatory diseases, we conducted Gene Ontology (GO) analysis of the 48 overlapping genes, which revealed enhanced interferon signaling and an inflammatory response in both RA and COVID-19 patients (Fig. 1, F and G). Specifically, the 259 RA-specific IFN-γ signature genes were linked to functions such as leukocyte activation and response to lipopolysaccharides, while the 31 COVID-19-specific genes were associated with cholesterol metabolism (Fig. 1, F and G). In addition, we found that the 170 genes that were downregulated in both RA synovial macrophages and IFN-γ-primed macrophages were related to small molecule transport and lipid localization (fig. S1, G and H). However, downregulated genes in COVID-19 monocytes did not show any significant enrichment in biological functions (fig. S1, G and H).

To further investigate the effects of cytokine stimulation on the transcriptomic signature of macrophages at the single-cell level, we analyzed single-cell RNA sequencing (scRNA-seq) data (GSE168710 (*18*)) from human monocyte-derived macrophages (HMDMs). Our findings indicated that HMDMs treated with both IFN-γ and TNF exhibited the most pronounced transcriptional changes, and these changes were most similar to those observed in HMDMs treated with IFN-γ alone (Fig. 1H). By comparing the HMDMs stimulated with each cytokine to control HMDMs, we identified genes regulated by IFN-γ at the single-cell level. Our analysis demonstrated that the IFN-γ-stimulated macrophage populations were most strongly associated with upregulated genes in patients with RA and acute COVID-19 (Fig. 1I and fig. S1I). Notably, some of the IFN-γ signature genes, including *EPSTI1, FCGR1A, GBP1, LAP3, STAT1, TNFSF10, UBE2L6*, and *VAMP5*, showed positive correlations with both RA and acute COVID-19 in the transcriptome of IFN-γ-stimulated macrophages (Fig. 1, F and J). Our results suggest that inflammatory diseases such as RA and COVID-19 exhibit a unique transcriptional landscape that is associated with specific IFN-γ signatures in human macrophages.

### IFN-γ signature of macrophages in heterogeneous inflammatory tissue environments

To gain insight into the heterogeneity within inflamed tissues, we analyzed scRNA-seq data of macrophages in bronchoalveolar lavage fluid (BALF) obtained from COVID-19 patients (GSE145926 (*41*)) and synovial fluid from RA patients (E-MTAB-8322 (*14*)) (Fig. 2A). In addition, we examined scRNA-seq data (GSE159117 (*42*)) from peripheral blood mononuclear cells (PBMCs) in RA patients to profile circulating monocytes in the blood. Using scRNA-seq data, we identified macrophage populations in BALF from healthy donors, patients with mild COVID-19, and severe COVID-19, and further characterized these populations based on their gene expression patterns (fig. S2A). UMAP analysis of scRNA-seq data revealed that macrophages were a significant portion of the BALF cell population. We identified five heterogeneous macrophage subpopulations including the MP-a (*S100A8* ^high^ *IFIT1* ^high^), MP-b (*APOE* ^high^ *FABP* ^high^ *CCL18* ^low^), MP-c (*APOE* ^high^), MP-d (*CCL18* ^high^), and MP-e (*S100A8* ^high^ *HSPA6* ^high^) subpopulations (fig. S2B). These macrophage subpopulations in BALF showed distinct distributions between healthy donors, mild and severe COVID-19 patients (Fig. 2B). Specifically, severe COVID-19 patients exhibited a significantly higher proportion of MP-a and MP-e subpopulations compared to healthy donors, while MP-b and MP-d had higher proportions in healthy donors (Fig. 2C). Next, we evaluated the IFN-γ signatures by comparing genes highly expressed in MP-a and MP-e with upregulated genes in IFN-γ-primed macrophages (fig. S2C). We identified 155 IFN-γ signature genes that are highly expressed in MP-a and MP-e. Expression of these signature genes was significantly increased in MP-a and MP-e compared to in MP-b, MP-c, and MP-d (fig. S2D). We also found that high expression of IFN-γ signature genes, such as *S100A8, S100A9, IFTM1, TNFSF10, VAMP5, GBP1,* and *GBP5*, was enhanced in patients depending on disease severity of COVID-19 patients (Fig. 2D). These results suggest that the increase of subpopulation of macrophages with distinct IFN-γ signatures in BALF is associated with the severity of COVID-19 and support the pathophysiological relevance of IFN-γ-mediated macrophages in severe COVID-19 patients.

**Fig. 2.**
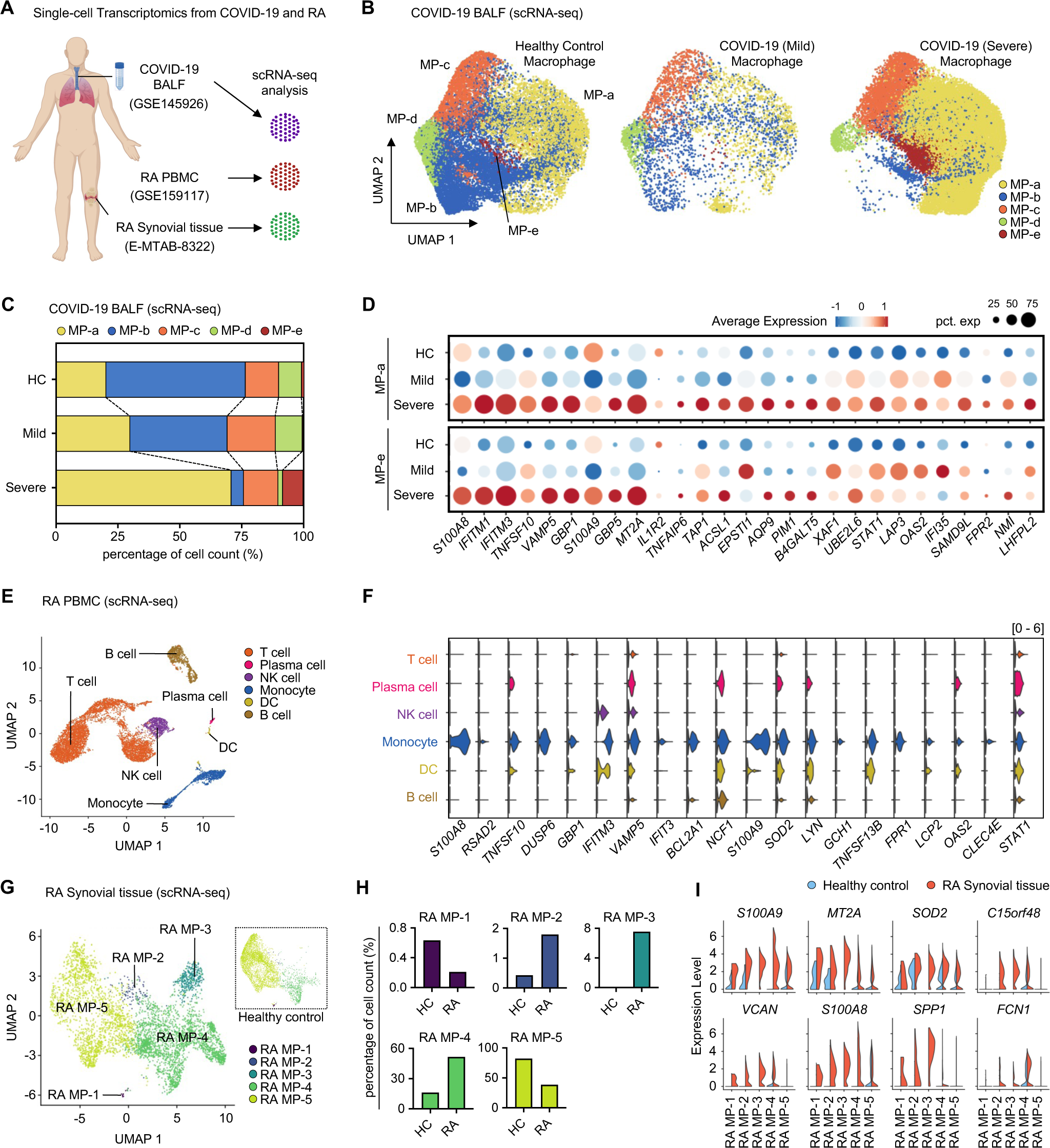
IFN-γ signature in macrophages of RA and COVID-19 patients. **(A)** Single-cell analysis overview. Single-cell RNA-seq data were acquired from GSE145926, GSE159117, and E-MTAB-8322, including bronchoalveolar lavage fluid (BALF) and peripheral blood mononuclear cells (PBMC). **(B)** UMAP plots depict five macrophage groups for controls (n = 4) and patients with moderate (n = 3) and severe (n = 6) COVID-19. **(C)** Bar plots illustrate macrophage cluster proportions, separated by condition and COVID-19 severity status. **(D)** Dot plots display the mean expression of IFN-γ signature-related genes for MP-a and MP-e clusters in controls (n = 4) and patients with moderate (n = 3) and severe (n = 6) COVID-19. The color range represents mean expression levels, while circle sizes indicate percentage expression. **(E)** Single-cell RNA-seq analysis of major cell types in PBMC of RA patients, using data from GSE159117. **(F)** Violin plot of IFN-γ signature gene expression in RA patients’ PBMCs. **(G)** UMAP plots present macrophage clusters from healthy controls and naïve RA patients in synovial tissue, with single-cell RNA-seq data from E-MTAB-8322. **(H)** Bar plots display macrophage cluster proportions for healthy controls and naïve RA patients in synovial tissue. **(I)** Violin plots show differentially expressed (DE) genes (FDR adjusted P < 0.05, fold change > 1.5) identified in synovial macrophage clusters from naïve RA patients, including both increasing and decreasing macrophage clusters.

To characterize the transcriptome of circulating monocytes in RA patients at the single-cell level, we conducted an scRNA-seq analysis of PBMCs obtained from RA patients (Fig. 2E). Our analysis revealed the presence of a CD14^+^ monocyte population within RA PBMCs (fig. S2E). We identified 134 differentially expressed genes in RA CD14^+^ monocytes that overlapped with upregulated genes in IFN-γ-primed macrophages (Fig. 2F and fig. S2F), indicating the presence of a distinct IFN-γ signature in circulating monocytes of RA patients prior to their infiltration into synovial tissue. Furthermore, we performed scRNA-seq analysis of macrophages in RA synovial tissue and classified them into five subpopulations based on unbiased gene expression patterns (Fig. 2G). Our analysis revealed that the MP-2, MP-3, and MP-4 subpopulations were increased in RA synovial tissue compared to healthy controls (Fig. 2H). These macrophage subpopulations showed high expression of IFN-γ signature genes, such as *S100A9*, *MT2A*, *SPP1* (*14*), and *SOD2*, involved in functions such as inflammation and defense responses (Fig. 2I and fig. S2G). Notably, similar gene expression patterns were observed in MP-a and MP-e from macrophages obtained from bronchoalveolar lavage fluid (BALF) in COVID-19 patients (Fig. 2I and fig. S2, H and I). Taken together, our single-cell transcriptome analyses provide evidence for the presence of macrophage heterogeneity and distinct IFN-γ signatures in inflamed tissues from both RA and COVID-19.

### Selective regulation of genes by JAK inhibitor in IFN-γ-primed macrophages

We investigated the regulatory mechanism of JAKi in IFN-γ-primed macrophages to comprehend the distinct IFN-γ signatures present in macrophages in RA and severe COVID-19. Our findings showed significantly higher levels of total STAT1 and STAT3 proteins in IFN-γ-primed macrophages compared to resting macrophages, along with elevated levels of phosphorylated-STAT1 and -STAT3 proteins (Fig. 3A and fig. S3A). We also observed that total protein levels of STAT1 and STAT3 in IFN-γ-primed macrophages increased over time, while the levels of phosphorylated-STAT1 and - STAT3 proteins peaked during the early stages of IFN-γ-priming at 3 h. Subsequently, we examined the ability of JAKi to modulate gene expression in IFN-γ-primed macrophages. We found that JAKi dose-dependently decreased the expression of the target genes of the IFN-γ-JAK-STAT axis, with the most significant inhibition observed at a JAKi concentration of 1 μM (Fig. 3B and fig. S3B). Moreover, we found that activated-STAT1 and -STAT3 in IFN-γ-primed macrophages were dephosphorylated rapidly to the level of resting macrophages within 1 h of JAKi treatment (Fig. 3B). We observed a gradual decrease in the total protein amount of STAT1 and STAT3, with the most pronounced decrease observed at 6 h (Fig. 3B). Finally, we investigated the temporal dynamics of JAKi-mediated inhibition of IFN-γ-induced genes. We found that JAKi suppressed the expression of *CCL2* and *CXCL10* in a time-dependent manner, with the greatest inhibition observed 6 h after JAKi treatment compared to 1 h after JAKi treatment (Fig. 3C). Our data suggest that JAKi efficiently inhibits the JAK-STAT pathway in IFN-γ-primed macrophages in vitro.

**Fig. 3.**
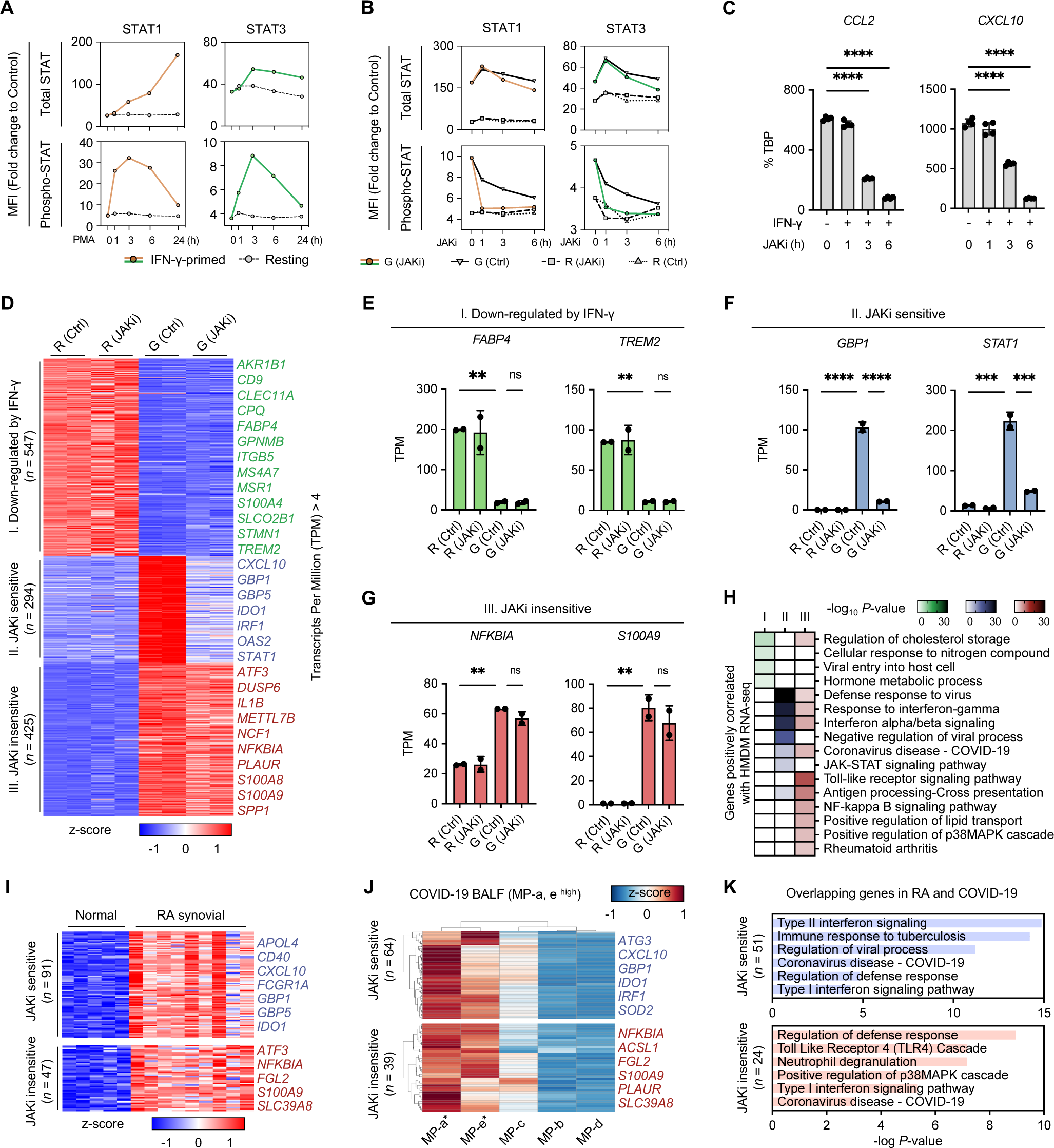
Distinct responses of IFN-γ-induced genes to JAK inhibitor. **(A)** Flow cytometry assessment of STAT1 and STAT3 protein levels in THP-1 cells treated with PMA and IFN-γ (or without) at various time points. Kinetics of total STAT and phospho-STAT tyrosine (STAT1, Tyr701; STAT3, Tyr705) measured by flow cytometry in IFN-γ-primed macrophages compared to resting macrophages. **(B)** Kinetics of total STAT1 (or STAT3) and phospho-STAT1 (or STAT3) measured by flow cytometry in JAK inhibitor-treated macrophages compared to resting and JAK inhibitor-untreated macrophages. Resting and IFN-γ-primed macrophages were differentiated for 24 h and then treated with JAKi (or DMSO) for up to 6 h. (A and B) Mean fluorescence intensity (MFI) represents a fold change compared to unstained samples. **(C)** RT-qPCR analysis of normalized target mRNA relative to TBP mRNA in THP-1 monocyte-derived macrophages under indicated conditions. IFN-γ-primed macrophages were treated with JAK inhibitor at a concentration of 1 μM for up to 6 h. Data show means ± SD from two independent experiments. **(D)** K-means clustering of differentially expressed (DE) genes in pairwise comparisons between the four conditions. DE genes identified by EdgeR (FDR adjusted P < 0.05, fold change > 2) were used. TPM values of RNA-seq data were filtered to be greater than 4. Non-significant clusters between replications were removed, resulting in three identified clusters. Clusters are indicated on the left. **(E-G)** Examples of expression for selected genes from clusters identified in the heatmap. Each dot on the bar plot represents one sample, and error bars denote the standard deviation. Error bars represent means ± SD. **(H)** Gene ontology (GO) analysis of THP-1 RNA-seq using genes positively correlated with HMDM RNA-seq. Heatmap displays the P-value (-Log10) significance of GO term enrichment for genes in each cluster, with clusters shown at the top. Downregulated by IFN-γ, n = 42; JAKi-sensitive, n = 129; JAKi-insensitive, n = 112. **(I)** Identification of genes associated with JAKi-sensitive or JAKi-insensitive in RA patients. Clusters are indicated on the left. **(J)** Identification of genes associated with JAKi-sensitive or JAKi-insensitive in COVID-19 patients. Clusters are indicated on the left. For heatmaps of single cells, mean expression values were used, and hierarchical analysis was performed. **(K)** GO analysis of overlapping JAKi-sensitive and JAKi-insensitive genes in RA or COVID-19 patients. Clusters are indicated on the left. JAKi-sensitive, n = 51; JAKi-insensitive, n = 24. p < 0.05(*), p < 0.01(**), p < 0.001(***) and p < 0.0001(****) by one-way ANOVA. GO analysis was performed using Metascape (http://metascape.org/).

To comprehensively understand the impact of JAKi on IFN-γ-primed macrophages, we conducted RNA-seq experiments and identified 1,709 differentially expressed (DE) genes through pairwise comparisons among the four conditions (fig. S3, C-E). Our findings revealed that JAKi caused more significant transcriptomic changes in IFN-γ-primed macrophages in comparison to resting macrophages (fig. S3, D and E). We further classified the total DE genes into three clusters based on *k*-means clustering (Fig. 3D). Cluster I (547 genes) contained IFN-γ-suppressed genes that remained unaffected by JAKi (Fig. 3, D and E). Notably, IFN-γ-induced genes were segregated into two clusters, cluster II (294 genes) and cluster III (425 genes), based on their JAKi sensitivity. The JAKi-sensitive genes in cluster II were related to interferon signaling and JAK-STAT pathways, which included *GBP1* and *STAT1* (Fig. 3, D, F, and H). In contrast, the JAKi-insensitive genes in cluster III were associated with lipid transport and MAPK cascade related functions, which included inflammatory genes such as *NFKBIA*, *PLAUR*, and *S100A9* (Fig. 3, D, G, and H). Both the JAKi-sensitive and JAKi-insensitive clusters exhibited enhanced defense responses to viruses and interferon-associated genes (Fig. 3H). To validate our findings, we compared our transcriptome data with publicly available data in human primary macrophages, which demonstrated similar regulation patterns by JAKi in IFN-γ signature genes (fig. S3, F and G).

Furthermore, we investigated whether JAKi sensitivity could divide IFN-γ signature genes in RA synovial macrophages and COVID-19 BALF macrophages (fig. S3H). Analysis showed JAKi-sensitive (*n* = 91) and JAKi-insensitive (*n* = 47) genes in RA synovial macrophages (Fig. 3I), and JAKi-sensitive (*n* = 64) and JAKi-insensitive (*n* = 39) genes in the MP-a and MP-e populations, which were highly represented in the BALF macrophages of COVID-19 patients (Fig. 3J). GO analysis indicated that JAKi-sensitive genes (*n* = 51) shared by RA and COVID-19 had enhanced interferon signaling and regulation of viral processes, while JAKi-insensitive genes (*n* = 24) shared by RA and COVID-19 enhanced toll-like receptor and MAPK cascade functions (Fig. 3K). Our findings suggest two distinct patterns of JAKi responsiveness in IFN-γ signature genes of macrophages, and the identified JAKi-sensitive and JAKi-insensitive genes are associated with RA and COVID-19, respectively. These selective differences offer insights into the molecular mechanisms that underlie how JAKi regulates inflammatory macrophages.

### Differential open chromatin changes by JAKi in IFN-γ-primed macrophages

To gain insight into the epigenetic mechanisms that contribute to the differential sensitivity of genes to JAK inhibition, we performed ATAC-seq under the same conditions as RNA-seq (fig. S3C). The quality of each ATAC-seq sample was evaluated based on fragment size (fig. S4A). We focused our analysis on the 9,461 differentially accessible chromatin regions (ACRs) (FDR < 0.05, |Fold change| > 2) identified in pairwise comparisons between the four conditions. Principal component analysis (PCA) plots of these differential peaks revealed a pattern similar to RNA-seq (fig. S4B). We identified three major ACR clusters, and the first cluster, comprising 3,470 ACRs closed by IFN-γ, showed a decrease in chromatin accessibility independent of JAKi, as demonstrated by the promoter and enhancers of the representative gene *CPQ* (Fig. 4, A and B). The second cluster, named “JAKi-sensitive” (*n* = 559), consisted of open chromatin regions that displayed *cis*-regulatory regions of the gene *GBP5*, which were induced by IFN-γ but significantly inhibited by JAKi (Fig. 4, A and B). In contrast, in the ACR cluster “JAKi-insensitive” (*n* = 5,432), open chromatin regions induced by IFN-γ were not affected by JAKi, as demonstrated in the enhancer region of the *S100A9* gene. To validate these findings, we confirmed that ATAC-seq data of HMDMs also showed JAKi-sensitive and JAKi-insensitive ACRs, as identified in THP-1-derived macrophages (fig. S4, C and D). Our results indicate that the responsiveness to JAKi is closely linked to open chromatin changes at the *cis*-regulatory elements of IFN-γ-induced genes in macrophages.

**Fig. 4.**
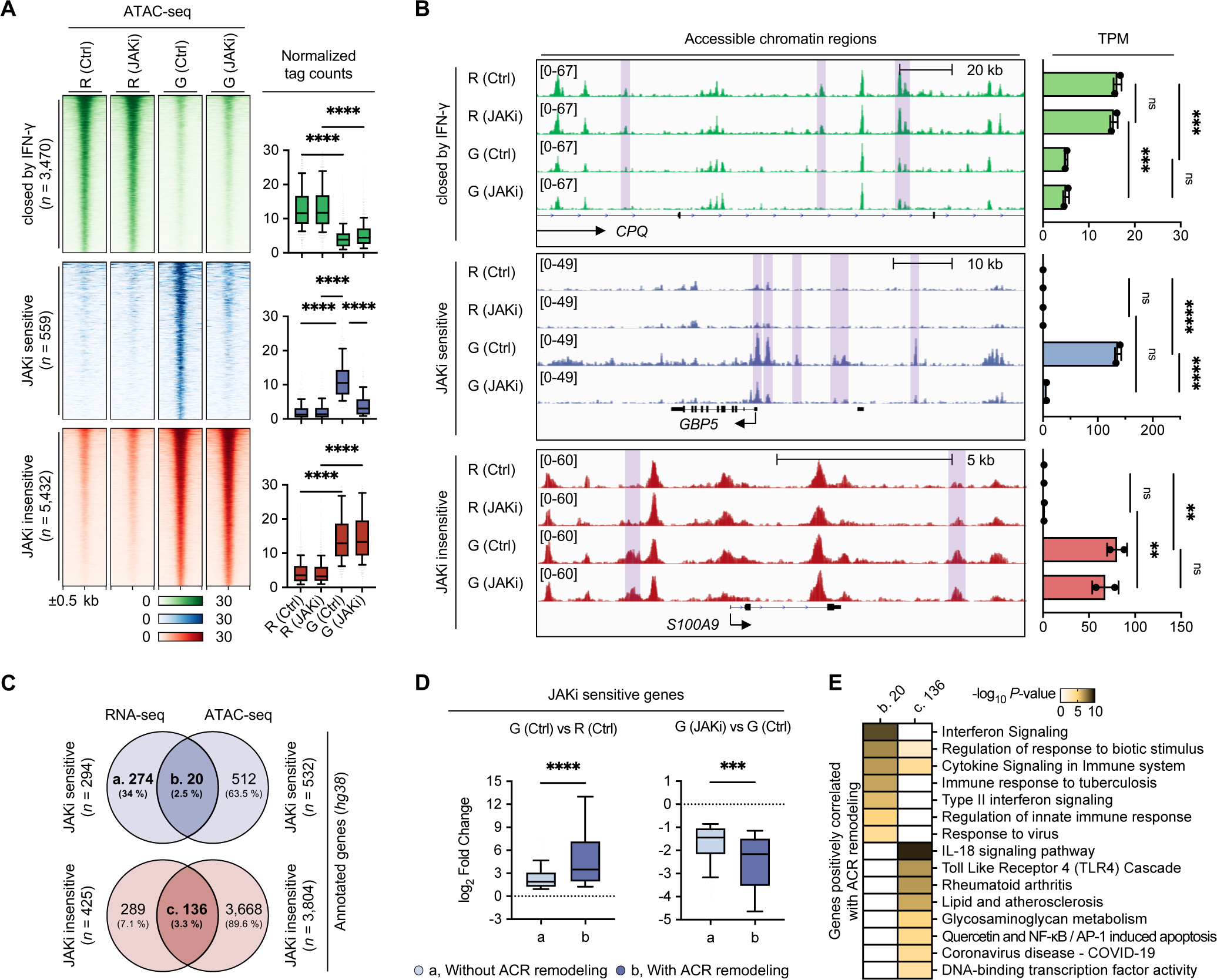
Differential sensitivity of chromatin accessibility to JAK inhibitor. **(A)** Heatmap of ATAC-seq signals in THP-1 macrophages with or without IFN-γ priming. Resting and IFN-γ-primed macrophages included JAKi treated and untreated conditions. Differentially accessible peaks (DP) were identified through edgeR (FDR adjusted P < 0.05, fold change > 2). Clusters are indicated on the left. Box plots represent normalized tag counts for ATAC peaks (right). **(B)** Example of ATAC-seq signals in a selected region from the cluster identified in the heatmap. IGV Genome browser tracks ATAC-seq signals in the vicinities of the indicated genes. Clusters are indicated on the left. Bar graphs display the expression levels of the indicated genes (right). Each dot on the bar plot represents one sample, and error bars denote the standard deviation. **(C)** Venn diagrams for identifying positively correlated genes in RNA-seq and ATAC-seq. For ATAC-seq analysis, an annotated gene in the hg38 genome was used. JAKi-sensitive ATAC-seq peaks annotated 532 genes. The JAKi-insensitive ATAC-seq peak annotated 3,804 genes. Annotation analysis was performed using HOMER. **(D)** Box plots illustrate the difference in fold change for genes with and without chromatin remodeling. JAKi-sensitive gene sets ’a’ and ’b’ identified by ATAC-seq in (C) were used. Fold changes in gene expression in IFN-γ-primed macrophages compared to resting macrophages were utilized. **(E)** GO analysis using genes identified in the integrated analysis of RNA-seq and ATAC-seq. Heatmap displays the P-value (-Log10) significance of GO term enrichment for each gene set, with gene sets shown at the top. b, JAKi-sensitive, n = 20; c, JAKi-insensitive, n = 136. Boxes encompass the twenty-fifth to seventy-fifth percentile changes. Whiskers extend to the tenth and ninetieth percentiles. The central horizontal bar represents the median. Error bars represent means ± SD. p < 0.05(*), p < 0.01(**), p < 0.001(***) and p < 0.0001(****) by one-way ANOVA. GO analysis was performed using Metascape (http://metascape.org/).

To further investigate the genes associated with differential ACRs, we performed an integrated analysis of RNA-seq and ATAC-seq. Our analysis revealed that 20 genes of the 294 JAKi-sensitive genes overlapped with annotated genes in ACR cluster “JAKi-sensitive,” while 136 of 425 JAKi-insensitive genes overlapped with annotated genes in ACR cluster “JAKi-insensitive” (Fig. 4C). The genes with ACR remodeling showed greater changes in response to IFN-γ and JAKi compared with genes without ACR remodeling (Fig. 4D). Functionally, JAKi-sensitive genes with ACR remodeling exhibited enhanced functions, such as interferon signaling and response to viruses (Fig. 4E). In contrast, JAKi-insensitive genes with ACR remodeling had enhanced functions, such as TLR4 cascade, lipid metabolism, and diseases, including RA, atherosclerosis, and COVID-19 (Fig. 4E). Overall, our findings suggest that IFN-γ-primed macrophages contain two distinct ACRs depending on JAKi sensitivity, and that these ACRs are involved in the regulation of IFN-γ-induced genes in macrophages.

### Distinct transcription factor repertoires in JAKi-regulated genes

We focused our investigation on the regulation of transcription factors associated with ACRs in IFN-γ-primed macrophages. To gain insights into the transcription factors predicted to be linked with ACRs regulated differentially by JAKi sensitivity, we examined the gene expression levels of transcription factors. We selected transcription factors differentially expressed for JAKi sensitivity from the three identified clusters in Fig. 3D (Fig. 5A). The JAKi-sensitive transcription factors (*n* = 34), including *EZH2*, *IRF1*, *IRF5*, *STAT1*, and *STAT3*, were repressed by JAKi in IFN-γ-primed macrophages (Fig. 5, A and B). On the other hand, the JAKi-insensitive transcription factors (*n* = 35), such as *CEBPB*, *ETS2*, *FOS*, *JUNB*, and *NFKBIA*, were not affected by JAKi in these macrophages (Fig. 5, A and C). Several AP-1 family members were detected in the transcription factor cluster for JAKi-insensitive genes (Fig. 5, A and C). We also investigated the differentially regulated transcription factors in RA synovial macrophages and COVID-19 BALF macrophages in Figs. 1 and 2. We observed that genes upregulated in RA synovial macrophages included JAKi-sensitive (*n* = 16) and JAKi-insensitive (*n* = 7) transcription factors (Fig. 5D). Similarly, genes upregulated in MP-a and MP-e of COVID-19 BALF also included JAKi-sensitive (*n* = 12) and JAKi-insensitive (*n* = 6) transcription factors (Fig. 5E). Intriguingly, JAKi-insensitive transcription factors such as *FOS*, *JUNB*, and *NFKBIA* were upregulated in both RA and COVID-19.

**Fig. 5.**
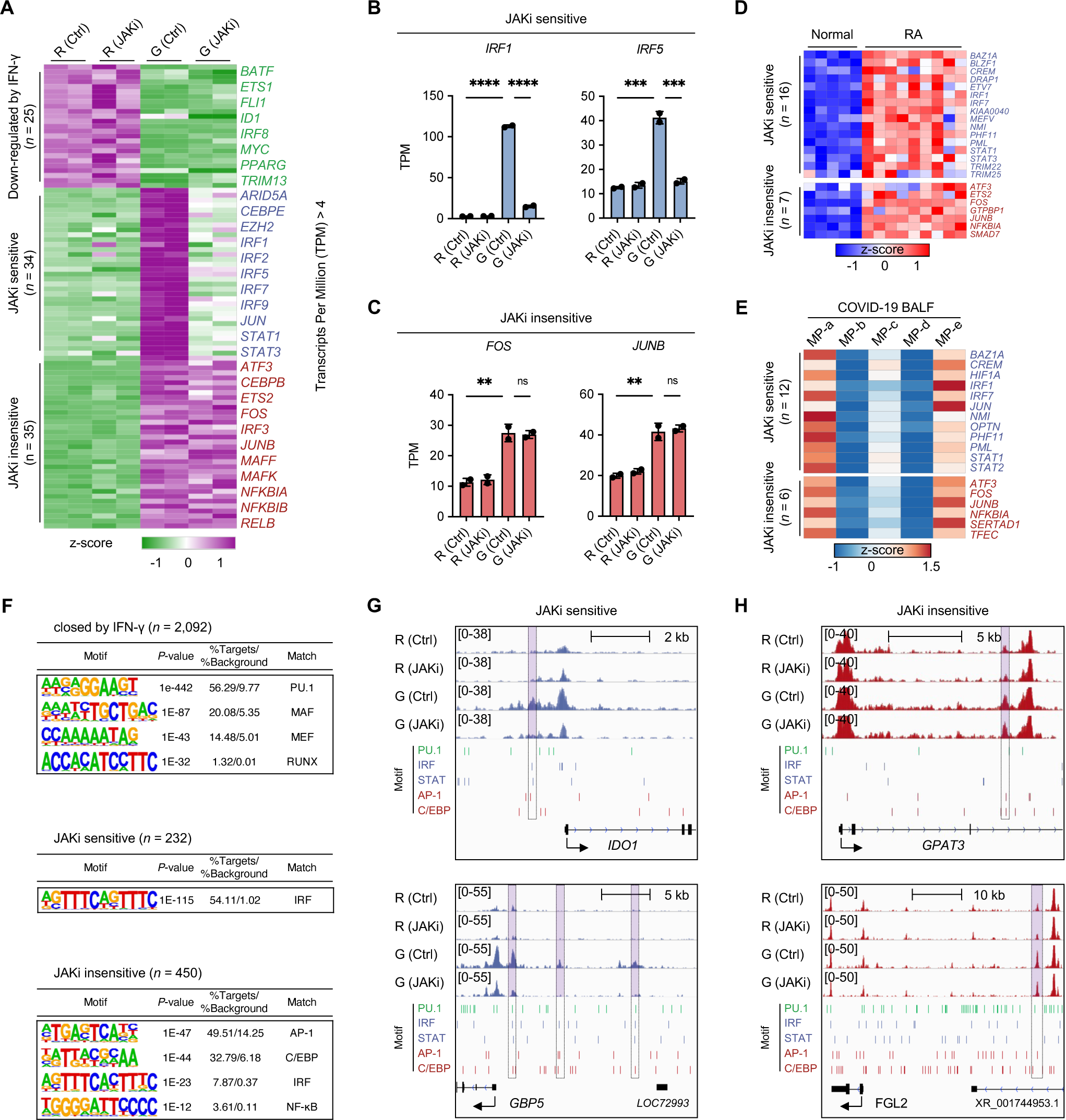
Divergent regulation of transcription factor repertoires in response to JAK inhibitor. **(A)** Heatmap of gene expression for 94 transcription factors (TF) in clusters defined by RNA-seq data analysis of THP-1 monocyte-derived IFN-γ-primed macrophages. Clusters are indicated on the left. **(B and C)** Examples of the expression of selected genes from JAKi-sensitive and JAKi-insensitive clusters identified in the heatmap for transcription factors. Each point on the bar plot represents one sample, and error bars denote the standard deviation. Error bars represent means ± SD. p < 0.05(*), p < 0.01(**), p < 0.001(***) and p < 0.0001(****) by one-way ANOVA. **(D)** Identification of transcription factor genes associated with JAKi-sensitive or JAKi-insensitive in patients with RA. Clusters are indicated on the left. **(E)** Identification of transcription factor genes associated with JAKi-sensitive or JAKi-insensitive in patients with COVID-19. Clusters are indicated on the left. Macrophage populations are indicated at the top. For single-cell heatmaps, mean expression values were used. **(F)** The most significantly enriched transcription factor (TF) motifs were identified by de novo motif analysis using HOMER in each cluster. Clusters are indicated at the top of each table. Peaks positively correlated with HMDM in THP-1 macrophages with and without IFN-γ priming were used. **(G and H)** The IGV Genome Browser displays known motifs in the vicinities of indicated genes for JAKi-sensitive (G) or JAKi-insensitive (H) chromatin regions. Clusters are indicated at the top. Known motif regions are denoted by colored lines. For PU.1, IRF, AP-1, and C/EBP motifs, the most significantly enhanced TF motifs identified in the bubble plot were used. PU.1 Motif, green; IRF and STAT Motif, blue; AP-1 and C/EBP Motif, red.

To gain further insights into the underlying mechanisms of transcription factors in the identified ACRs, we performed *de novo* motif analysis (Fig. 5F). Our analysis revealed that the cluster of ACRs “closed by IFN-γ” was enriched for PU.1 and MAF motifs, which are consistent with previously reported negative IFN-γ signatures (*21*). The “JAKi-sensitive” cluster showed significant enrichment in IRF motifs, similar to the study of STAT1 and IRF1 in IFN-γ-primed macrophages (*43*). In contrast, the JAKi-insensitive cluster showed enhanced motifs for AP-1 and C/EBP. The AP-1 protein family acts as a crucial transcription node, which integrates inputs from the upstream MAPK signaling pathway (*44*). The enhanced AP-1 motif in the JAKi-insensitive ACR cluster of IFN-γ-primed macrophages is consistent with the enhanced function of the “p38MAPK cascade” in the JAKi-insensitive gene cluster identified in Fig. 3.

To validate the results of *de novo* motif analysis, we further investigated the known motifs for JAKi-sensitive and JAKi-insensitive ACRs. Our analysis revealed distinct patterns of PU.1, IRF, and AP-1 motifs for each ACR cluster (fig. S5A). To confirm these results, we analyzed the motif density of PU.1, IRF, STAT, AP-1, and CEBP in the representative gene locus of each ACR cluster. We observed that the PU.1 motif was significantly enriched in the *cis*-regulatory regions of “closed by IFN-γ” cluster genes, such as *ITGB5, SLCO2B1*, and *LRMP* (fig. S5B). We also found that JAKi-sensitive ACRs were enriched with IRF and STAT motifs at the promoter or enhancer regions of JAKi-sensitive genes, including *IDO1* and *GBP5* (Fig. 5G and fig. S5C). In contrast, ACRs of the “JAKi-insensitive” cluster showed only AP-1 and CEBP motifs in the open chromatin regions of genes, such as *GPAT3* and *FGL2* (Fig. 5H and fig. S5D), instead of IRF and STAT motifs. Collectively, our results suggest that the distinct repertoire of transcription factors for ACRs identified in IFN-γ-primed macrophages might be associated with differential regulation of JAKi sensitivity.

### JAKi selectively suppresses STAT1/IRF1-bound *cis*-regulatory regions

To explore the impact of JAKi sensitivity on transcription factor binding to each ACR cluster, we analyzed ChIP-seq data from IFN-γ-primed macrophages. Our analysis revealed distinct STAT1 and STAT3 binding patterns across the three ACR clusters, with both “JAKi-sensitive” and “JAKi-insensitive” clusters showing increased binding in IFN-γ-primed macrophages relative to resting macrophages. As further support for the involvement of IRF1 in regulating JAKi-sensitive genes, IRF1 ChIP-seq signals were predominantly enriched in the “JAKi-sensitive” ACR cluster of IFN-γ-primed macrophages (Fig. 6A), consistent with the *de novo* motif analysis shown in Fig. 5. To compare the binding occupancy of STATs and IRF1 between “JAKi-sensitive” and “JAKi-insensitive” ACRs, we evaluated the normalized tag counts for each cluster. We found that binding signals of STAT1 and IRF1 were significantly higher in the “JAKi-sensitive” cluster, while STAT3 exhibited minimal differences between two clusters (Fig. 6C).

**Fig. 6.**
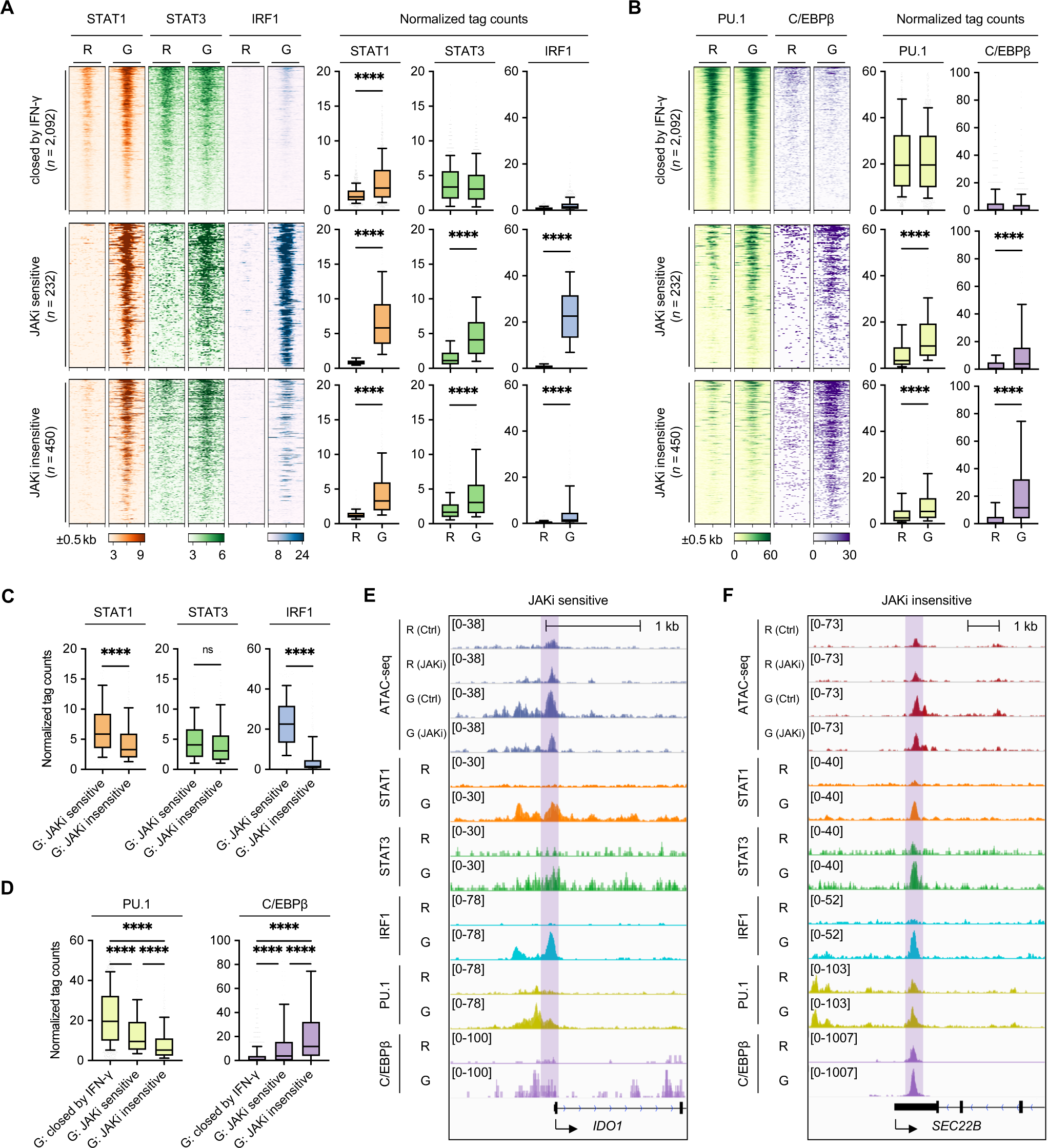
Distinct transcription factors bind to open chromatin regions regulated by JAKi. (A and B) Heatmap of ChIP-seq signals in human monocyte-derived macrophages (HMDM) with or without IFN-γ priming. ChIP-seq data from HMDM were obtained from GSE43036, GSE120943, and GSE98367. ChIP-seq signals were identified in ATAC-seq clusters of THP-1 monocyte-derived macrophages. Clusters are indicated on the left. Box plots represent normalized tag counts for ChIP-seq (right). **(C and D)** Boxplot comparing normalized tag counts for each cluster using IFN-γ-primed macrophage conditions. **(E and F)** The IGV genome browser displays ChIP-seq signals in the vicinities of genes marked as JAKi-sensitive (E) or JAKi-insensitive (F) chromatin regions. Clusters are indicated at the top. ChIP-seq and macrophage conditions are shown on the left. R, Resting macrophages; G, IFN-γ-primed macrophages. Boxes encompass the twenty-fifth to seventy-fifth percentile changes. Whiskers extend to the tenth and ninetieth percentiles. The central horizontal bar represents the median. Error bars represent means ± SD. p < 0.05(*), p < 0.01(**), p < 0.001(***) and p < 0.0001(****) by an unpaired t-test. R, Resting macrophages; G, IFN-γ-primed macrophages.

We also examined the binding preferences of other transcription factors. We found that PU.1 preferentially bound to the ACRs of the “closed IFN-γ” cluster, while C/EBPβ was most strongly bound to the ACRs of the “JAKi-insensitive” cluster in IFN-γ-primed macrophages (Fig. 6, B and D, and fig. S6, A-C). The representative gene track demonstrated preferential occupancy of IRF1 and STAT1 at the ACRs of JAKi-sensitive genes, such as *IDO1* and *GBP5* (Fig. 6E and fig. S6B). Conversely, C/EBPβ was found to strongly bind to *cis*-regulatory regions of JAKi-insensitive genes, such as *SEC22B* and *HCAR2* (Fig. 6F and fig. S6C). Taken together, our findings indicate that JAKi selectively inhibits IRF1/STAT1-bound open chromatin regions of JAKi-sensitive genes, while JAKi-insensitive ACRs preferentially interact with C/EBPβ and possibly with AP-1 transcription factors.

### JAKi-sensitivity of IFN-γ-signature genes in MTX-resistant RA and COVID-19 vaccinated donors

To explore the JAKi sensitivity of IFN-γ signature genes in RA patients who are resistant to MTX, we analyzed scRNA-seq data of synovial macrophages by comparing MTX-resistant RA patients with untreated naïve RA patients (Fig. 7A). We observed a higher frequency of two distinct macrophage subpopulations (MP-2 and MP-3) in MTX-resistant RA patients compared to naïve RA patients (Fig. 7B). To further investigate the JAKi sensitivity of IFN-γ signature genes, we analyzed highly expressed genes in MP-2 and MP-3 macrophage subpopulations (fig. S7A). We found that both MP-2 and MP-3 expressed JAKi-sensitive genes, including *GBP1* and *CCL2* (MP-2, *n* = 21; MP-3, *n* = 8), as well as JAKi-insensitive genes, such as *FOS* and *JUNB* (MP-2, *n* = 22; MP-3, *n* = 34) (Fig. 7C and fig. S7A).

**Fig. 7.**
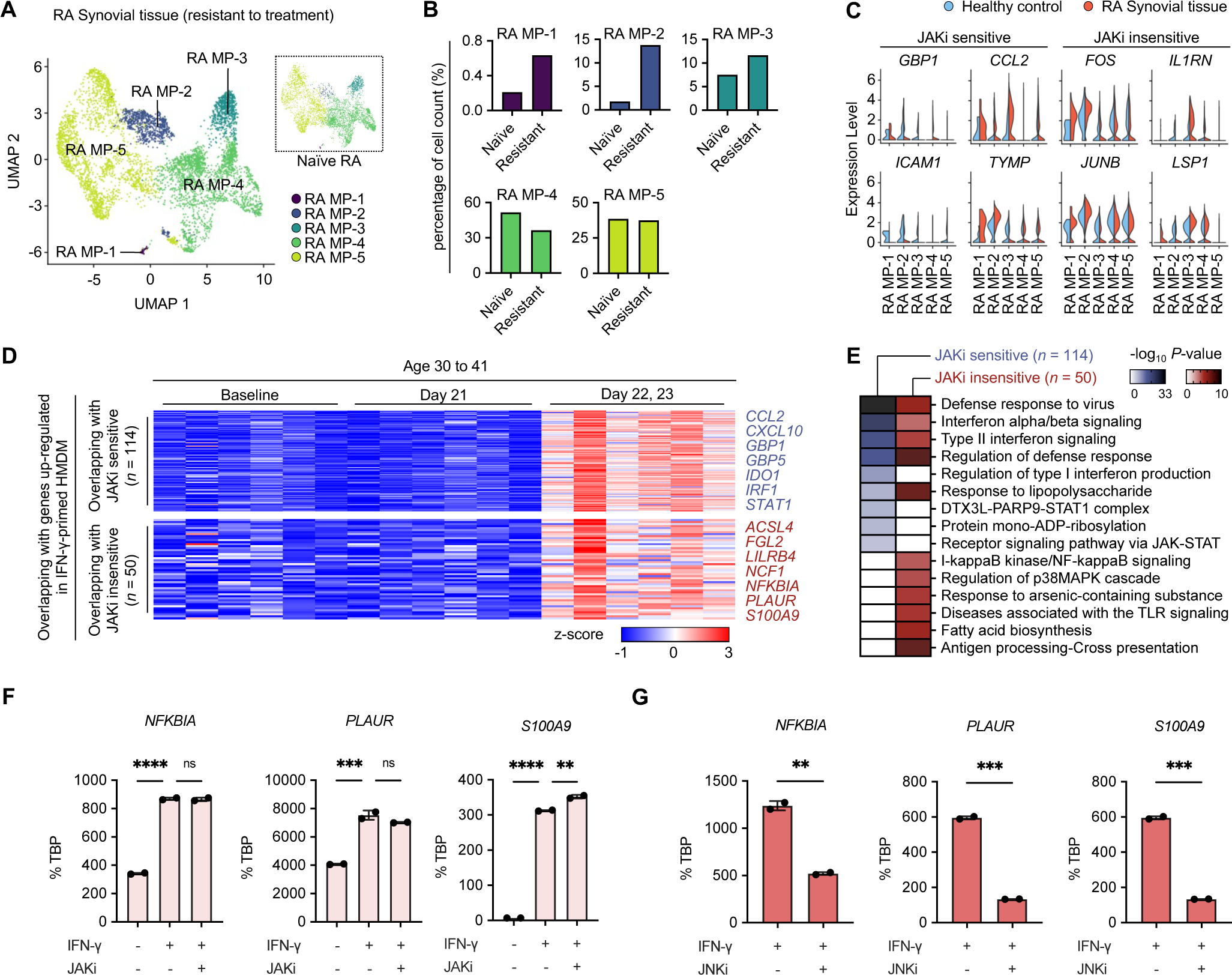
Sensitivity of IFN-γ signature genes to JAKi in methotrexate-resistant RA patients and COVID-19 vaccinated individuals. **(A)** Analysis of scRNA-seq data for macrophage types in synovial tissue of patients with RA. UMAP plots display macrophage clusters from drug-resistant RA patients compared with naïve RA patients. **(B)** Bar plots illustrate macrophage cluster proportions for naïve RA patients and drug-resistant RA patients. **(C)** Violin plots for genes identified in synovial macrophage clusters from drug-resistant RA patients. Differentially expressed (DE) genes (FDR adjusted P < 0.05, fold change > 1.5) identified in increasing macrophage clusters compared with decreasing macrophage clusters in the synovial tissue of drug-resistant RA patients were used. **(D)** Heatmap of gene expression after COVID-19 vaccination. Identified JAKi-sensitive and JAKi-insensitive genes were used. Clusters are shown on the left. **(E)** Gene Ontology (GO) analysis of genes associated with IFN-γ-primed macrophages in patients vaccinated against COVID-19. Genes from IFN-γ-primed macrophages that positively correlate with HMDM were used. Heatmap displays the P-value (-Log10) significance of GO term enrichment for genes in each cluster. GO analysis was performed using Metascape (http://metascape.org/). Overlapping with JAKi-sensitive, n = 114; Overlapping with JAKi-insensitive, n = 50. **(F)** RT-qPCR analysis of normalized target mRNA compared with TBP mRNA in THP-1 monocyte-derived macrophages under the indicated conditions. THP-1 monocyte-derived macrophages were induced with PMA (100 nM) and IFN-γ (100 U/ml) for 24 h. IFN-γ-primed macrophages were treated with JAK inhibitor (JAKi, tofacitinib) at a concentration of 1 μM for 6 h. **(G)** RT-qPCR analysis of normalized target mRNA compared with TBP mRNA in THP-1 monocyte-derived macrophages under the indicated conditions. THP-1 monocyte-derived macrophages were induced with PMA (100 nM), IFN-γ (100 U/ml), and JNK inhibitor (JNKi, SP600125, 10 μM) for 24 h. Data show means ± SD from two independent experiments. p < 0.05(*), p < 0.01(**), p < 0.001(***) and p < 0.0001(****) by one-way ANOVA.

To explore the relationship between JAKi-sensitivity of IFN-γ signature genes and COVID-19 vaccine efficacy, we conducted an analysis of RNA-seq data (GSE169159(*45*)) from whole blood samples of COVID-19 vaccinated donors. Our results showed that there were distinct DE gene patterns in samples of donors aged 30-41 years on day 22 or 23 post-vaccination (fig. S7B). Additionally, we selected six donors who exhibited the most significant transcriptional changes among the identified patients aged 30-41 years, and these samples were clearly clustered by the date of post-vaccination (fig. S7, C and D). We identified 1,285 upregulated genes related to the functions of defense responses to the virus, interferon signaling, and inflammatory responses, on days 22 and 23 compared to 21 days post-vaccination for the six selected donors (fig. S7, E and F). These upregulated genes by COVID-19 vaccination included both JAKi-sensitive (*n* = 114) genes, such as *GBP5* and *IDO1*, and JAKi-insensitive (*n* = 50) genes, such as *S100A9* and *NFKBIA* (Fig. 7D).

Recent research has suggested that compromised type I and II IFN immunity is responsible for the cessation of the immune response in age-related diseases, leading to increased disease severity (*46*). Notably, our study found that the upregulation of IFN-γ signature genes following COVID-19 vaccination was not significantly increased in donors aged 62-74 years (fig. S7G). Interestingly, we observed that JAKi-sensitive genes were associated with defense response to virus, interferon signaling, and JAK-STAT signaling pathway while the JAKi-insensitive genes linked to COVID-19 vaccination showed interferon-related functions, p38MAPK cascade, and antigen presentation (Fig. 7E). These findings imply that JAKi regulation of interferon signature genes might have an impact on the efficacy of COVID-19 vaccination.

We also investigated whether JAKi-insensitive genes could be targeted by JNK inhibitor (JNKi) in IFN-γ-primed macrophages, given that the functions of JAKi-insensitive genes exhibited MAPK pathways (Fig. 3, H and K). Our results revealed that treatment with JNKi significantly reduced the expression of JAKi-insensitive genes, such as *NFKBIA, PLAUR*, and *S100A9* (Fig. 7, F and G). These findings suggest that JNK inhibition represents an alternative strategy to suppress IFN-γ signature genes that are resistant to JAKi regulation. These findings suggest that the molecular mechanisms of JAKi-related regulation of IFN-γ signature genes are implicated in different aspects of both RA and COVID-19, including MTX resistance in RA patients and the efficacy of COVID-19 vaccination.

## Discussion

The interferon signature is a well-established hallmark of chronic inflammatory diseases, such as lupus and rheumatoid arthritis (*16, 23, 47*). In the inflammatory environment, interferon stimulation activates the JAK-STAT signaling pathway, and JAK inhibitors (JAKi) have been shown to effectively reduce abnormal inflammation by inhibiting this pathway (*48, 49*). However, the precise epigenetic mechanisms by which JAKi regulates gene expression in macrophages remain incompletely understood. Additionally, recent large-scale clinical studies have raised concerns about the long-term safety of JAKi treatment, and its potential association with cardiovascular diseases and cancer (*28, 37, 50, 51*). Therefore, it is crucial to investigate the molecular mechanisms underlying the interferon signature to comprehensively understand the benefits and risks of JAKi in various disease conditions. This study aimed to investigate the epigenetic mechanisms by which JAKi regulates IFN-γ signature genes in human macrophages.

Inflammatory cytokines are known to cause a range of responses in macrophages, resulting in a heterogeneous phenotype of pathogenic macrophages commonly found in inflamed tissues (*8, 12*). Among these cytokines, IFN-γ plays a prominent role in the polarization of M1-like inflammatory macrophages, which are frequently observed in chronic inflammatory diseases (*15, 16*). In this study, our comparative analysis revealed that IFN-γ signature genes exhibited not only uniquely induced genes but also commonly induced genes with other inflammatory stimuli, such as TNF and LPS. Importantly, we observed that these signature genes were predominantly present in patients with RA and COVID-19. Our single-cell transcriptome analysis revealed that IFN-γ signature genes were enriched in specific cell subpopulations associated with disease severity in both RA and COVID-19. These genes were present in RA monocytes and synovial macrophages, as well as COVID-19 BALF macrophages. These findings suggest that IFN-γ signature genes may play a crucial role in the pathogenesis of chronic inflammatory diseases.

The prolonged exposure of macrophages to cytokines can lead to memory-like changes, called ‘innate immune memory’ or ‘trained immunity’ (*52–54*). These changes are closely linked to epigenetic and metabolic modifications. The IFN-γ signature is particularly important in this context since it is associated with long-lasting transcriptomic and epigenomic changes that promote chronic inflammation and make it difficult to treat diseases. The resulting epigenomic signature from IFN-γ priming amplifies the transcriptional response of inflammatory cytokines, such as TNF and IL-6, while repressing homeostatic and anti-inflammatory gene expression programs (*39, 43*). Although JAK inhibitors primarily target the signaling pathway, our research demonstrates that JAKi-induced transcriptomic changes are also influenced by the epigenomic signatures in IFN-γ-primed macrophages. Our study reveals that JAKi effectively suppresses a subset of IFN-γ-induced genes that are associated with interferon signaling and defense response. However, some IFN-γ-induced genes related to TLR signaling and the MAPK cascade remain unresponsive to JAKi, despite showing STAT1 and STAT3 binding at the promoter or enhancers. The intrinsic difference between JAKi-sensitive and JAKi-insensitive genes can be attributed to distinct DNA binding motifs of *cis*-regulatory elements. Specifically, JAKi preferentially targets IRF-STAT motifs to inhibit JAKi-sensitive genes, whereas JAKi-insensitive genes contain AP1-C/EBP motifs. Recent studies have suggested that increased chromatin accessibility of IRF-enriched open chromatin regions over AP-1-enriched regions plays a critical role in the trained CD14^+^ monocytes and antiviral immunity of myeloid cells induced by influenza vaccination with adjuvant (*55, 56*). Our findings provide valuable insights into the underlying mechanisms of how JAKi selectively regulates IFN-γ-induced genes based on epigenomic landscapes.

The balance between STAT1 and STAT3 plays a crucial role in regulating inflammatory responses in macrophages, as these transcription factors have similar DNA binding motifs and can compete for binding to the same sites or form heterodimers or homodimers (*57*). While IFN-γ priming predominantly activates STAT1 over STAT3, our findings demonstrate that IFN-γ activates both STAT1 and STAT3. The binding preferences of partner transcription factors with STAT1 and STAT3 are determined by DNA binding motifs in distinct open chromatin regions. For example, IRF1, an essential binding partner of STAT1, regulates TLR responses by increasing chromatin accessibility in macrophages (*58*). Our ChIP-seq analysis revealed that IRF1 preferentially binds to open chromatin regions of JAKi-sensitive genes with STAT1 and STAT3. On the other hand, C/EBPβ, one of the lineage-determining transcription factors in macrophages, predominantly binds to JAKi-insensitive open chromatin regions with STAT1 and STAT3. Both IRF1 and C/EBPβ can act as potential pioneer factors in macrophages, suggesting that their differential binding preferences may contribute to the distinct changes in chromatin accessibility induced by JAK inhibition (*58, 59*). IFN-γ priming can activate autocrine loops mediated by chemokines or low levels of TNF and TNFSF members, resulting in the AP-1, C/EBP, and NF-kB motif enrichments observed in the JAKi-insensitive genes. Moreover, we found that among AP-1 family members, *JUNB* and *FOS* are JAKi-insensitive genes enriched with AP-1 motifs in open chromatin regions. To test the possibility of overcoming resistance to JAKi in IFN-γ-primed macrophages, we examined a JNK inhibitor in our experiments. Our results revealed that the JNK inhibitor suppresses some JAKi-insensitive genes, such as *NFKBIA, PLAUR*, and *S100A9*, indicating that it could be a potential alternative strategy to repress JAKi-insensitive genes.

JAK inhibitors have become an important treatment option for RA patients who are resistant to MTX (*49*). Our study utilizing single-cell transcriptome analysis revealed an increase in the frequency of certain macrophage subpopulations, specifically MP-1 and MP-2, in MTX-resistant RA patients compared to naïve RA patients. These subpopulations include both JAKi-sensitive and JAKi-insensitive IFN-γ signature genes. Interestingly, JAKi-sensitive genes, such as *CCL2* and *GBP1*, may be potential targets for JAKi therapy in MTX-resistant RA patients, while JAKi-insensitive genes, including *FOS* and *LSP1*, may limit the effectiveness of JAKi therapy and contribute to the safety of JAKi. Our transcriptomic analysis also showed an increase in IFN-γ signature genes, including both JAKi-sensitive and JAKi-insensitive genes, after 3 weeks of COVID-19 vaccination. However, these responses were not significant in older vaccinated subjects (*45*). The suppression of IFN-γ signature genes by JAKi, particularly in older people, would be interesting to investigate to understand the mechanisms behind lower vaccination efficacy in individuals associated with aging and immunosuppression. Taken together, drug resistance in RA and immune response following COVID-19 vaccination are associated with IFN-γ signature genes, including JAKi sensitivity (*60*). Therefore, further research is needed to fully elucidate the mechanisms underlying how JAK inhibitors differentially regulate IFN-γ signature genes across diverse clinical contexts of human diseases (*61–63*).

In summary, our findings shed light on the mechanisms underlying the epigenetic regulation of IFN-γ signature genes by JAKi in human macrophages. This study provides insights into the potential therapeutic benefits and risks of JAKi treatment for various chronic inflammatory diseases. Further research is warranted to fully understand the epigenetic regulation of the interferon signature and its implications for JAKi therapy. Ultimately, a deeper understanding of these mechanisms may pave the way for the development of more targeted and effective therapies for patients with chronic inflammatory diseases.

## Materials and Methods

### Cell Culture

Human THP-1 cells were obtained from Korean Cell Line Bank (KCLB40202) and were cultured in RPMI-1640 (Gibco™, 11875-119) supplemented with 10% Fetal Bovine Serum (Gibco, 16000-044) and 1% Penicillin/streptomycin (Gibco, 15070-063). The THP- 1 cell line was routinely cultured in a humidified 5% CO2 incubator at 37 °C. THP-1 monocytes were differentiated into macrophages by incubation with 100 ng/ml Phorbol 12-myristate 13-acetate (PMA) (Sigma-Aldrich, P1585) for 24 h. THP-1-monocyte-derived IFN-γ-primed macrophages were induced by co-stimulation with PMA and 100 U/ml human IFN-γ (Roche, 11040596001) for 24 h. Following priming, the macrophages were treated with JAK inhibitors (Tofacitinib) (Sigma-Aldrich, PZ0017-5MG) at up to 1 μM for a maximum of 6 h. JNK inhibitors (Medchemexpress, HY-12041) were treated at 10 μM on THP-1 monocytes with PMA and IFN-γ for 24 h.

### Quantitative Real-Time PCR

Total RNA was extracted from THP-1-monocyte-derived macrophages using RibospinTM II (GeneAll Biotechnology, 314-150), and 500 ng of total RNA was reverse transcribed using the RevertAid First Strand cDNA Synthesis kit (ThermoFisher™, K1622). Real-time PCR was performed with TOPreal™ qPCR 2X PreMIX (Enzynomics, RT500M) and Rotor-Gene Q software 2.3.1 (QIAGEN). Primer sequences are provided in the Table S2. RNA samples were stored at -80 °C to proceed with RNA sequencing.

### Omni-ATAC-seq Library Preparation

The Omni-ATAC-seq library was prepared as previously described (*64, 65*). THP-1-monocyte-derived macrophages of 50,000 cells cultured in a medium were pretreated with 200 U/ml Deoxyribonuclease I (Worthington, LS002004) at 37 °C for 30 min to remove free-floating DNA and digest DNA from dead cells. This medium was washed with Dulbecco’s phosphate buffered saline (BIOWEST, L0615-500). After washing, 50,000 cells of THP-1-monocyte-derived macrophages were harvested from the culture plate using TrypLE™ (Gibco™, 12604013). 50,000 viable cells were centrifuged at 500 g for 5 min at 4 °C in a fixed-angle centrifuge. The macrophage pellet was washed once more with cold ATAC-seq resuspension buffer (RSB; 10 mM Tris-HCl pH 7.5 (Invitrogen™, 15567-027), 10 mM NaCl (Invitrogen™, AM9759), and 3 mM MgCl2 (Invitrogen™, AM9530G) in Nuclease-Free Water (QIAGEN, 981103)). Cells were centrifuged at 500 g for 5 min at 4 °C in a fixed-angle centrifuge. After centrifugation, 900 μl of supernatant was aspirated, which left 100 μl of supernatant. This remaining 100 μl of supernatant was carefully aspirated by pipetting with a P200 pipette tip to avoid the cell pellet. Cell pellets were then resuspended in 50 μl of ATAC-seq RSB containing 0.1% NP40 (Roche, 11332473001), 0.1% Tween-20 (Roche, 11332465001), and 0.01% Digitonin (Promega, G9441) by pipetting up and down three times. This cell lysis reaction was incubated on ice for 3 min. After lysis, 1 ml of ATAC-seq RSB containing 0.1% Tween-20 (without NP40 or digitonin) was added, and the tubes were inverted to mix. Nuclei were then centrifuged for 10 min at 500 g in a pre-chilled (4 °C) fixed-angle centrifuge. The supernatant was removed with two pipetting steps, as described before, and nuclei were resuspended in 50 μl of transposition mix (25 μl TD Tagment DNA Buffer (Illumina, 15027866), 2.5 μl TDE1 Tagment DNA Enzyme (Illumina, 15027865), 16.5 μl DPBS, 0.5 μl 1% digitonin, 0.5 μl 10% Tween-20, and 5 μl Nuclease-Free Water) by pipetting up and down six times. Transposition reactions were incubated at 37 °C for 30 min in a thermomixer with shaking at 1,000 g. Reactions were cleaned up with MinElute Reaction Cleanup Kit (QIAGEN, 28206). Reactions were pre-amplified for 5 cycles using NEBNext® High-Fidelity 2X PCR Master Mix (NEB, M0541L) with adapter primers. Adapter primer sequences are provided in the Table S1. After PCR amplification, the amplification profile was manually evaluated, and the number of additional cycles required for amplification was determined. Buenrostro *et al*. 2015 were consulted to determine the additional cycles required for ATAC-seq library amplification (*65*). Final amplified library samples were stored at -80 °C for ATAC-seq after library purification with AMPure XP beads (Beckman, A63880).

### Flow Cytometry

Cultured 1 x 10^6^ THP-1 is immediately fixed by adding Fixation buffer (BioLegend, 420801) pre-warmed to 37 °C. Cells are fixed at 37 °C for 15 minutes. Fixed cells are centrifuged at 350 g for 5 minutes at room temperature. After centrifuging the cells, remove the supernatant and resuspend the pellet in Cell Staining Buffer (BioLegend, 420201). Centrifuge the resuspended cells at 350 g for 5 min at room temperature, then remove the supernatant and resuspend. Resuspend the cells by adding True-Phos™ Perm Buffer (BioLegend, 425401) pre-chilled to -20 °C and permeabilize for at least 60 minutes at - 20 °C. Permeabilized cells are centrifuged at 1000 g for 5 min at room temperature. After centrifuging the cells, remove the supernatant and resuspend the pellet. Resuspended cells are washed twice with Cell Staining Buffer. Resuspend the washed cells in Cell Staining Buffer. Transfer 2 x 10^5^ cells / 50 µl to a 1.5 ml tube. Add antibody (Alexa Fluor® 647 Mouse anti-Total Stat1, BD biosciences, 58560; PE anti-STAT1 Phospho (Tyr701), BioLegend, 666404; APC Mouse anti-Total Stat3, BD biosciences, 560392; PE/Cyanine5 anti-STAT3 Phospho (Tyr705), BioLegend, 651014) cocktail(s) to appropriate tubes, vortex to mix, and incubate for 30 minutes at room temperature in the dark. Add 1 ml of Cell Staining Buffer, centrifuge cells at 1000 g at room temperature for 5 minutes, decant supernatant. Repeat for a total of two washes. Resuspend cells in approximately 500 µl of Cell Staining Buffer and analyze with a flow cytometer (CytoFLEX, Beckman Coulter). The data were analyzed using FlowJo™ (v10.8.1) software.

### Genome Alignment and Annotations

All human data are aligned and annotated against the *hg38* reference genome. Reads were aligned to the *hg38* genome using the *hg38*.fa (https://hgdownload.soe.ucsc.edu/goldenPath/hg38/bigZips/hg38.fa.gz) and *hg38*.gtf (from the table browser on the USCS website) files provided by USCS. To annotate aligned reads, we used “*hg38*.ncbiRefSeq.gtf” files provided by USCS. (https://hgdownload.soe.ucsc.edu/goldenPath/hg38/bigZips/genes/hg38.ncbiRefSeq.gtf.gz).

### Data Mapping

The raw sequencing data were quality filtered using Trim Galore (v0.6.6). Preprocessed reads were mapped to the human *hg38* genome. RNA-seq data were mapped to the *hg38* genome using STAR aligner (v2.7.3a) with default parameters (*66*). ChIP-seq data were mapped to the *hg38* genome using Bowtie2 (v2.3.5.1) with default parameters (*67*). ATAC-seq data were mapped to the *hg38* genome using Bowtie2 with the parameters *”--very-sensitive --no-discordant -X 2000*” (https://github.com/harvardinformatics/ATAC-seq, last updated: 2019). HOMER (v4.11.1) was used to convert aligned reads into ‘‘Tag Directory’’ for further analysis (*68*).

### Bulk RNA-seq Analysis

The RNA-seq data used in this study consisted of publicly available data obtained from the Gene Expression Omnibus (GEO) under accession numbers GSE98368 (*21*), GSE120944 (*39*), GSE100382 (*38*), GSE169159 (*45*), GSE198256 (*40*), and the data generated from our own experiments. Each mapped read was quantified using the ‘‘*analyzeRepeats*’’ script of HOMER. To generate a table of raw read counts, the parameters *-count exons -condenseGenes -noadj* were used. To generate a table of TPM values, the parameters *-count exons -condenseGenes -tpm* were used. Differentially expressed (DE) genes were identified using raw sequencing read counts by edgeR (v3.30.3) analysis via the ‘‘*getDifferentialExpression*’’ HOMER command at *P*-adj (adjusted *P*-value) < 0.05 and FC (Fold Change) > 2 (*69*). We removed genes with a TPM value of less than 4 (or 2) from the identified DE genes. Gene ontology analysis was performed using Metascape (*70*). Venny (v2.1.0) and Circos (v0.69-8) were performed to identify genes showing positive correlations between RNA-seq data (*71, 72*).

### ChIP-seq and ATAC-seq Analysis

The ChIP-seq and ATAC-seq data used in this study consisted of published data obtained from GEO under accession numbers GSE43036 (*43*), GSE120943 (*39*), and GSE98365 (*21*), and the data generated from our own experiments. For ChIP-seq and ATAC-seq analysis, preprocessed reads were aligned to reference the human genome using bowtie2 with parameters. The quality check of ATAC-seq was performed with ATACseqQC (v1.12.5) (*73*). We used the “*makeTagDirectory*” followed by “*findPeaks*” command from HOMER to identify peaks of ChIP-seq enrichment over the background. A false discovery rate (FDR) threshold of 0.001 was used for all data sets. ChIP-seq and ATAC-seq peaks were used with 2-fold changes compared to controls. The bedgraph file was created using the “*makeUCSCfile*” HOMER command, with the “*-fragLength given - o auto -fsize 5e7 -res 1*” parameter. The bigwig file was created using the “*bamCoverage*” deepTools (v3.5.0) command, with the “*--binSize 10 --normalizeUsing RPGC -- effectiveGenomeSize 2913022398*” parameter (*74*). ATAC-seq of THP-1 monocyte-derived IFN-γ-primed macrophages identified significant clusters using overlapping peaks for each condition. ACR regions regulated by JAKi in IFN-γ primed macrophages were excluded when identifying “JAKi insensitive” clusters.

### Single-cell (sc) RNA-seq Analysis

The pre-processed gene-cell matrices of the single-cell RNA-seq data were downloaded from the GEO or EMBL’s European Bioinformatics Institute (EMBL-EBI) database (accession number GSE168710 (*18*), GSE145926 (*41*), GSE159117 (*42*) and E-MTAB-8322 (*14*)). These matrices were subsequently loaded into Seurat (v4.1.0) R package for quality control, filtering, and downstream analyses (*75*). Cells with a mitochondrial gene percentage greater than 10%, or fewer than 200 genes or more than 6,000 genes, were filtered out. Some samples were cut off more strongly according to the reference paper. The gene expression data was then normalized using a log normalization method, and principal component analysis (PCA) was performed to reduce the dimensionality of the data. The significant principal components were selected based on the “elbow plot” method, and clustering was performed using the shared nearest neighbor (SNN) algorithm. Differentially expressed (DE) genes between different clusters were identified using the Wilcoxon rank-sum test. Cell type identification is based on known cell type markers and was further confirmed through reference articles. The UMAP plot was generated to visualize the cell clusters and the differentially expressed genes.

### Microarray Data Analysis

The microarray data used in this study used public data obtained from GEO under accession number GSE97779 (*21*). Microarray data analysis was performed using the Gene Expression Omnibus 2R (GEO2R) online tool. Differential expression analysis was performed between control macrophages and RA synovial macrophages using GEO2R. DE genes with FDR < 0.05 and fold change > 2 were considered significant.

### Motif Enrichment

HOMER’s motif analysis (*findMotifsGenome.pl*), including known default motifs and de novo motifs, was used to identify motifs enriched in differentially regulated ACRs. Significant motifs identified through known motifs analysis were screened in genomic tracks with ACRs.

### Data Visualization

The Integrative Genomics Viewer (IGV, v2.9.2) and deepTools (v3.5.0) were used to visualize ChIP-seq and ATAC-seq data (*76*). RNA-seq data were visualized with heatmap using Morpheus (http://software.broadinstitute.org/morpheus).

### Statistical Analysis

Statistical tests were selected based on appropriate assumptions with respect to data distribution and variance characteristics. Statistical significance was defined as p < 0.05. The whiskers of boxplots represent the 10-90th percentiles of the data. Statistical analyses were performed using GraphPad Prism 9 (GraphPad Software, v9.3.1).

## General

We thank Dr. Lionel Ivashkiv for helpful discussions and review of the manuscript.

## Author contributions

G.K. conceptualized, designed, and performed most of the experiments and performed bioinformatic analysis. G.K. and Y.P. contributed experiments and expertise. Y.P. and Keunsoo K. performed bioinformatic analysis. Kyuho. K. conceptualized and oversaw the study and edited the manuscript. All authors reviewed and provided input on the manuscript.

## Data and materials availability

The data sets generated by the authors will be deposited in the GEO database. Reviewers can request the access token to the data. This study did not generate new unique reagents.

