## Supplementary Materials for "Selective epigenetic regulation of IFN-γ signature genes by JAK inhibitor in inflammatory diseases"

Geunho Kwon *et al.*

**This PDF file includes:**

Fig. S1. Identification of distinct IFN-γ signatures of monocytes and macrophages in inflammatory diseases.

Fig. S2. Identification of IFN-γ signatures in tissue macrophages from RA and COVID-19.

Fig. S3. Identification of IFN-γ-induced genes differentially regulated by JAKi in RA and COVID-19.

Fig. S4. Identification of distinct chromatin remodeling for JAKi in IFN-γ-primed macrophages.

Fig. S5. Identification of differential motif landscapes for regions of chromatin remodeling regulated by JAKi.

Fig. S6. Binding of identified TFs to open chromatin regions regulated by JAKi.

Fig. S7. Identification of the JAKi-insensitive signature gene in inflammatory diseases.

Table S1. Illumina/Nextera i5 common adapter and i7 index adapters.

Table S2. Primers for RT-qPCR.

### Supplementary Figure 1

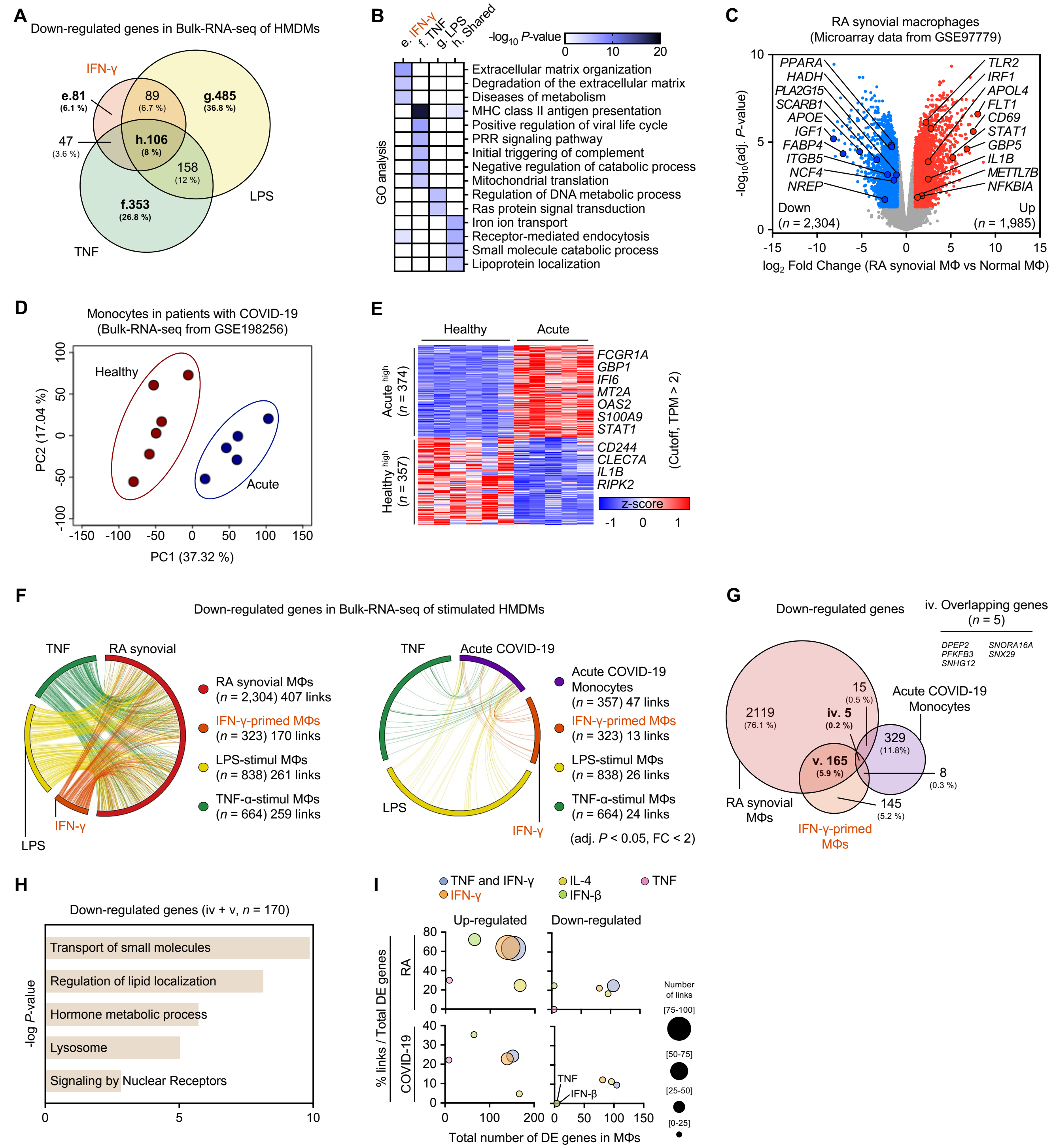

**Fig. S1. Identification of distinct IFN-γ signatures of monocytes and macrophages in inflammatory diseases.**

**(A)** Venn diagrams for down-regulated genes identified in stimulated HMDMs. RNA-seq data of human monocyte-derived macrophages (HMDM) from GSE98368, GSE120944, and GSE100382 were used. Down-regulated genes identified by EdgeR (FDR adjusted $P < 0.05$, fold change < 2) compared with resting macrophages were used. TPM values of RNA-seq data were cut off to be greater than 4. IFN-γ, orange; TNF, green; LPS, yellow. **(B)** Gene ontology (GO) analysis using genes specifically down-regulated or shared in each stimulated HMDM RNA-seq. The 'e', 'f', 'g' and 'h' genes identified in (A) were used. e, IFN-γ, $n = 81$; f, TNF, $n = 353$; g, LPS, $n = 485$; h, Shared, $n = 106$. **(C)** Volcano plot of transcriptional changes between healthy controls and synovial macrophages in patients with RA. Colored dots correspond to genes with significant (FDR adjusted $P < 0.05$) and greater than two-fold expression changes. Microarray data were obtained from GSE97779 (Normal macrophages, $n = 5$; RA synovial macrophages, $n = 9$). **(D)** Principal component analysis (PCA) plots of RNA-seq samples. TPM values for genes ($n = 10{,}000$) identified in RNA-seq analysis was used. For monocyte RNA-seq from patients with COVID-19, healthy monocyte samples ($n = 6$) and acute COVID-19 monocyte samples ($n = 5$) that did not interfere with each other were used in the PCA plot. Each point represents an RNA-Seq sample. Samples with similar gene expression profiles are clustered together. Bulk RNA-seq were obtained from GSE198256 (Healthy : GSM5942339, GSM5942340, GSM5942341, GSM5942343, GSM5942348, GSM5942349; Acute COVID-19 : GSM5942352, GSM5942353, GSM5942354, GSM5942355, GSM5942356;). **(E)** Heatmap for differentially expressed (DE) genes in acute COVID-19 compared with healthy conditions. DE genes identified by EdgeR (FDR adjusted $P < 0.05$, fold change > 2) were used. TPM values of RNA-seq data were cut off to be greater than 2. Clusters are indicated on the left. **(F)** Circos plots display down-regulated genes linked between RA synovial macrophages (or acute COVID-19 monocytes) and human macrophages primed or stimulated by cytokines. Links are colored according to priming or stimulation. **(G)** Venn diagram of genes down-regulated in RA synovial macrophages, acute COVID-19 monocytes and IFN-γ-primed HMDM. Genes shared by all data sets are indicated on the right. **(H)** GO analysis using down-regulated genes shared by RA synovial macrophages, acute COVID-19 monocytes, and IFN-γ-primed HMDM. The 'iv' and 'v' genes identified in (G) were used. Bar plot showing the *P*-value (-Log10) significance of GO term enrichment for genes in each cluster. iv, $n = 5$; v, $n = 165$. **(I)** Bubble plot analyzing the ratio of DE gene link numbers with RA synovial macrophages or acute COVID-19 monocytes compared with the DE gene of each human monocyte-derived stimulated macrophage. The dot size represents the number of links. GO analysis was performed using Metascape (http://metascape.org/).

### Supplementary Figure 2

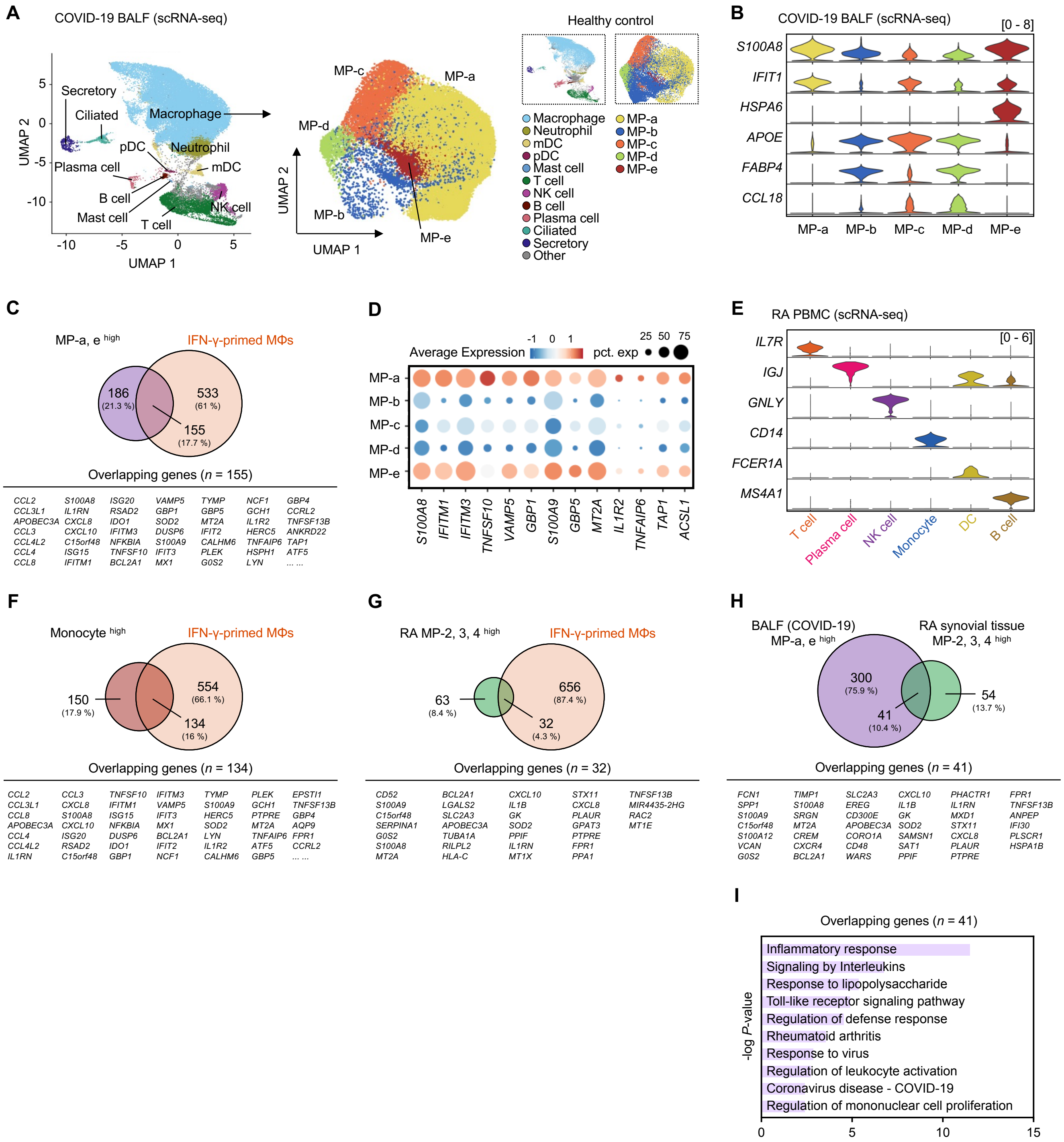

**Fig. S2. Identification of IFN-γ signatures in tissue macrophages from RA and COVID-19.**

**(A)** Analysis of scRNA-seq data for major cell types in BALFs (*n* = 13). single-cell RNA-seq data were obtained from GSE145926. Macrophage populations were extracted separately and re-clustered. Clusters are colored and labeled in UMAP space. UMAP plots for healthy controls are shown on the right. **(B)** Violin plot of gene expression for macrophage clusters in BALF. BALF macrophage clusters were classified into five groups according to the indicated gene expression levels. **(C)** Venn diagrams for up-regulated genes in both IFN-γ-primed macrophages and BALF. In scRNA-seq of BALF, DE genes (FDR adjusted $P < 0.05$, fold change > 1.5) for an increasing macrophage population in the patient were identified compared with a decreasing macrophage population in the patient. Genes shared by the data sets are shown below. **(D)** Dot plot of mean expression of IFN-γ signature related genes for each macrophage cluster, as indicated. Color ranges indicate mean expression levels and circle sizes indicate percentage expression. **(E)** Violin plots for marker genes representing cell types identified in PBMCs of patients with RA. Cell-type marker genes correspond to genes with more than 1.5-fold significant (FDR adjusted $P < 0.05$) expression changes compared with other clusters. **(F)** Venn diagrams for up-regulated genes in both IFN-γ-primed macrophage and monocyte clusters of RA PBMC. In scRNA-seq of PBMCs, DE genes (FDR adjusted $P < 0.05$, fold change > 1.5) for monocytes were identified compared with other cell types. Genes shared by the data sets are shown below. **(G)** Venn diagrams for up-regulated genes in both IFN-γ-primed macrophages and naïve RA synovial tissue. In scRNA-seq of synovial tissue, DE genes (FDR adjusted $P < 0.05$, fold change > 1.5) for an increasing macrophage population in the patient were identified compared with a decreasing macrophage population in the patient. Genes shared by the data sets are shown below. **(H)** Venn diagrams for up-regulated genes in both BALF and naïve RA synovial tissue. Genes shared by the data sets are shown below. **(I)** Gene ontology (GO) analysis using overlapping genes in RA synovial macrophages and COVID-19 BALF macrophages. GO analysis was performed using Metascape (http://metascape.org/). Overlapping genes, *n* = 41.

**Supplementary Figure 3**

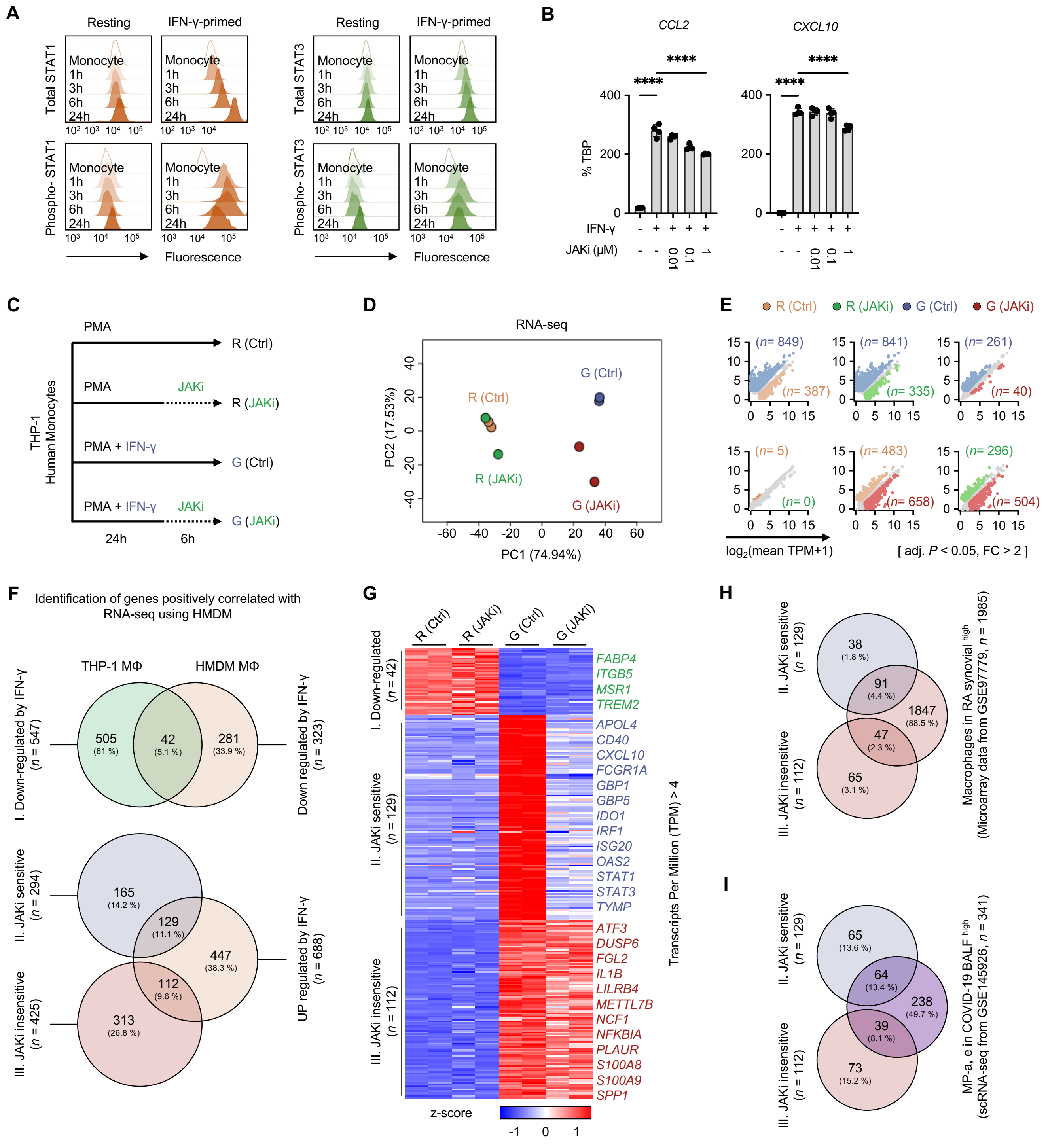

**Fig. S3. Identification of IFN-γ-induced genes differentially regulated by JAKi in RA and COVID-19.**

**(A)** Histograms of STAT1 and STAT3 protein levels in THP-1 cells treated with PMA and IFN-γ (or without) at different times. Histograms show the fluorescence intensity for each time point. **(B)** RT-qPCR analysis of normalized target mRNA compared with TBP mRNA in THP-1 monocyte-derived macrophages under the indicated conditions. THP-1 monocyte-derived macrophages were induced with PMA (100 nM) and IFN-γ (100 U/ml) for 24 h. IFN-γ-primed macrophages were treated with JAK inhibitor (JAKi, tofacitinib) at concentrations of up to 1 μM for 1 h. Data show means ± SD from two independent experiments. $p < 0.05$(*), $p < 0.01$(**), $p < 0.001$(***) and $p < 0.0001$(****) by one-way ANOVA. **(C)** Experimental design: Treated with PMA (Phorbol 12-myristate 13-acetate, 100 nM) for THP-1 monocyte-induced macrophage differentiation. THP-1 monocyte-derived macrophages were primed or not with IFN-γ (100 U/ml) for 24 h. Resting or IFN-γ-primed macrophages were treated or not with JAKi (tofacitinib, 1 μM) for 6 h. Harvested cells were used for RNA-seq and ATAC-seq. R(Ctrl), Without priming and treatment; R(JAKi), treatment with JAKi without priming; G (Ctrl), primed with IFN-γ without JAKi treatment; G (JAKi), Primed with IFN-γ and treated with JAKi. **(D)** Principal component analysis results of RNA-seq samples. Total differentially expressed ($n$ = 1,709) genes identified by edgeR (FDR adjusted $P < 0.05$, fold change > 2) for all pairwise were used. TPM values of RNA-seq data were cut off to be greater than 4. Each point represents an RNA-Seq sample. Samples with similar gene expression profiles are clustered together. **(E)** Scatter plot showing the DE gene for each condition. Total DE genes identified by EdgeR (FDR adjusted $P < 0.05$, fold change > 2) were used ($n$ = 1,709). Conditions are indicated on either the x-axis or the y-axis. Dots for the DE gene were conditionally colored. R (Ctrl), orange; R (JAKi), green; G (Ctrl), blue; G (JAKi), red. **(F)** Venn diagrams to identify positively correlated genes in THP-1 monocyte-derived IFN-γ-primed macrophages and human monocyte-derived IFN-γ-primed macrophages. RNA-seq data of human monocyte-derived macrophages (HMDM) from GSE98368 were used. Differentially expressed genes identified by edgeR (FDR adjusted $P < 0.05$, fold change > 2) in HMDM are used. TPM values of HMDM RNA-seq data were cut off to be greater than 4. **(G)** Heatmap of THP-1 RNA-seq using genes positively correlated with HMDM RNA-seq. Clusters are indicated at the left. **(H and I)** Venn diagram to identify genes associated with IFN-γ-primed macrophages in patients with RA or COVID-19. Genes from IFN-γ-primed macrophages that positively correlate with HMDM were used.

**Supplementary Figure 4**

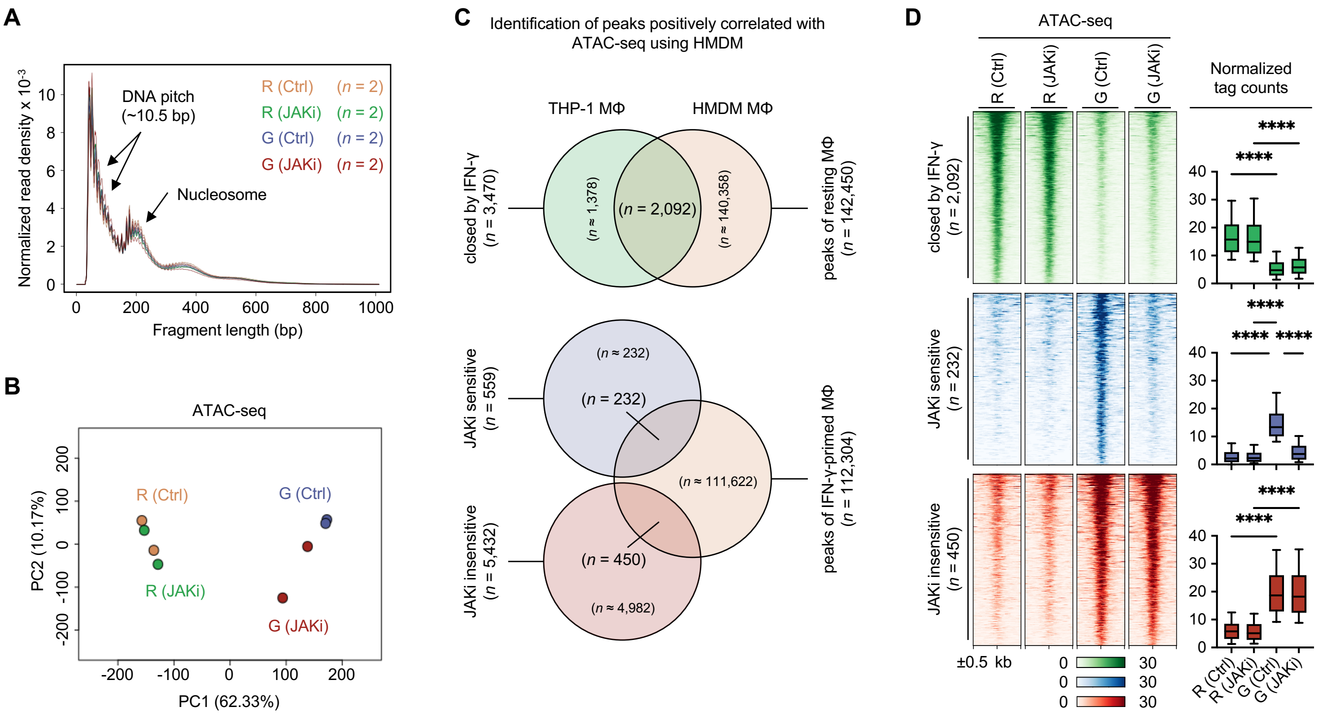

**Fig. S4. Identification of distinct chromatin remodeling for JAKi in IFN-γ-primed macrophages.**

**(A)** ATAC-seq fragment sizes generated from THP-1 nuclei indicate chromatin-dependent periodicity with a spatial frequency consistent with nucleosomes and a high-frequency periodicity consistent with the pitch of the DNA helix for fragments less than 200 bp. Fragment length indicates the genomic distance between two Tn5 insertion sites, as determined by paired-end sequencing of ATAC fragments. Density indicates the fraction of fragments with the indicated length. R (Ctrl), orange; R (JAKi), green; G (Ctrl), blue; G (JAKi), red. **(B)** Principal component analysis results of ATAC-seq (Assay for Transposase-Accessible Chromatin using sequencing) samples. Total differential peaks (DP) identified by edgeR (FDR adjusted $P < 0.05$, fold change > 2) for all pairwise were used ($n = 38{,}950$). Normalized tag count values of DP were used. Each point represents an ATAC-Seq sample. Samples with similar peak profiles were clustered together. **(C)** Venn diagrams to identify positively correlated peaks in THP-1 monocyte-derived IFN-γ-primed macrophages and human monocyte-derived IFN-γ-primed macrophages. ATAC-seq data of human monocyte-derived macrophages (HMDM) from GSE98365 were used. Differentially peaks identified by edgeR (FDR adjusted $P < 0.05$, fold change > 2) in HMDM are used. **(D)** Heatmap of THP-1 ATAC-seq using peaks positively correlated with HMDM ATAC-seq. Clusters are indicated at the left. Box plots represent normalized tag counts for ATAC peaks (right). Boxes encompass the twenty-fifth to seventy-fifth percentile changes. Whiskers extend to the tenth and ninetieth percentiles. The central horizontal bar represents the median. Error bars represent means ± SD. $p < 0.05$(*), $p < 0.01$(**), $p < 0.001$(***) and $p < 0.0001$(****) by one-way ANOVA.

### Supplementary Figure 5

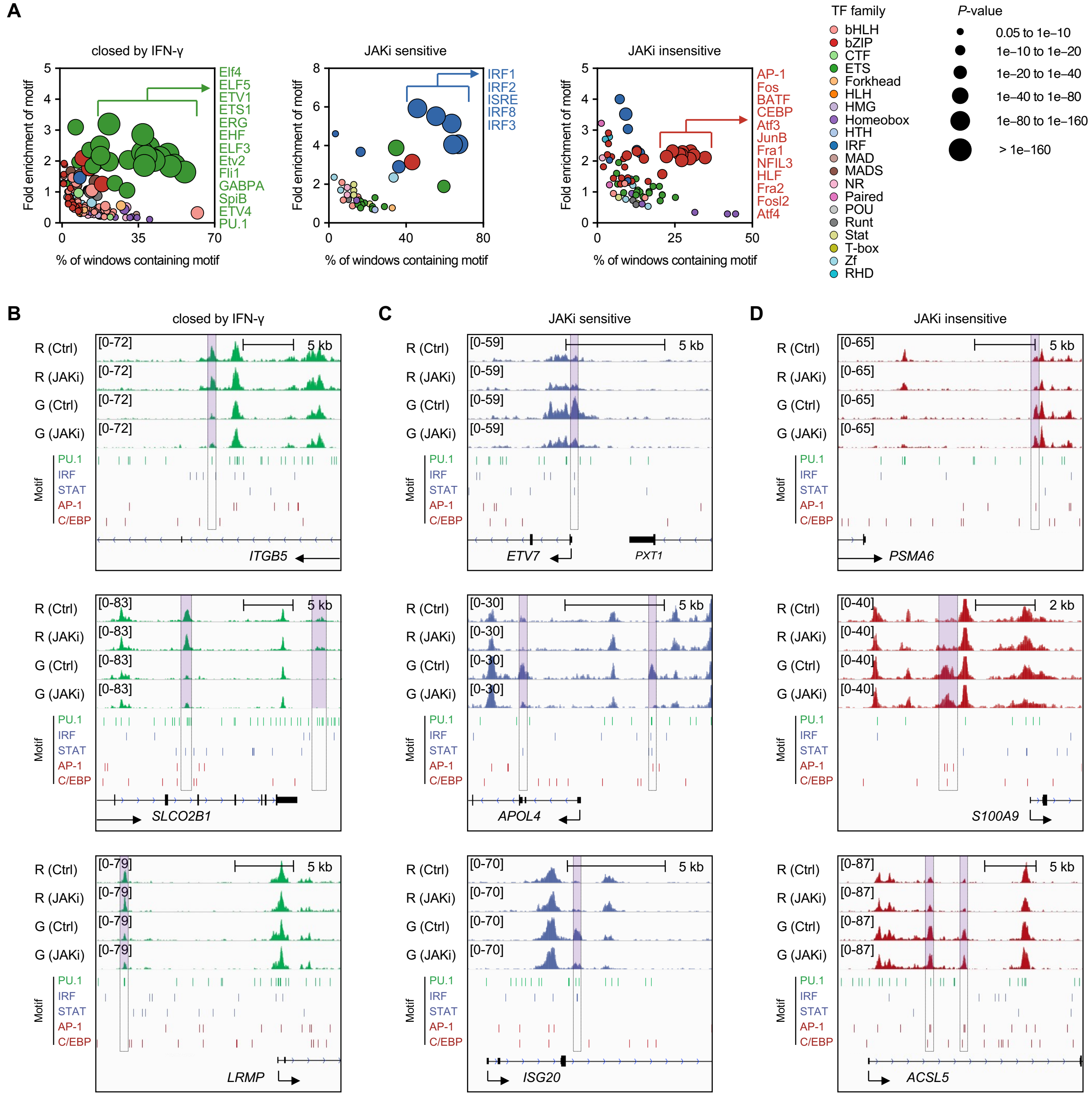

**Fig. S5. Identification of differential motif landscapes for regions of chromatin remodeling regulated by JAKi.**

**(A)** Bubble plot of transcription factor (TF) motif enrichment analysis for each cluster. The motifs were identified by known motif analysis using HOMER. Clusters are indicated at the top. The most significantly enriched TF motif is shown on the right. Color range depicts different transcription factor families and circle size refers to *P*-value significance. **(B-D)** The IGV Genome Browser displays known motifs in the vicinities of indicated genes for chromatin remodeling regions. Clusters are indicated at the top. Known motif regions are indicated by colored lines. For PU.1, IRF, AP-1 and C/EBP motifs, the most significantly enhanced TF motifs identified in the bubble plot were used. PU.1 Motif, green; IRF and STAT Motif, blue; AP-1 and C/EBP Motif, red.

**Supplementary Figure 6**

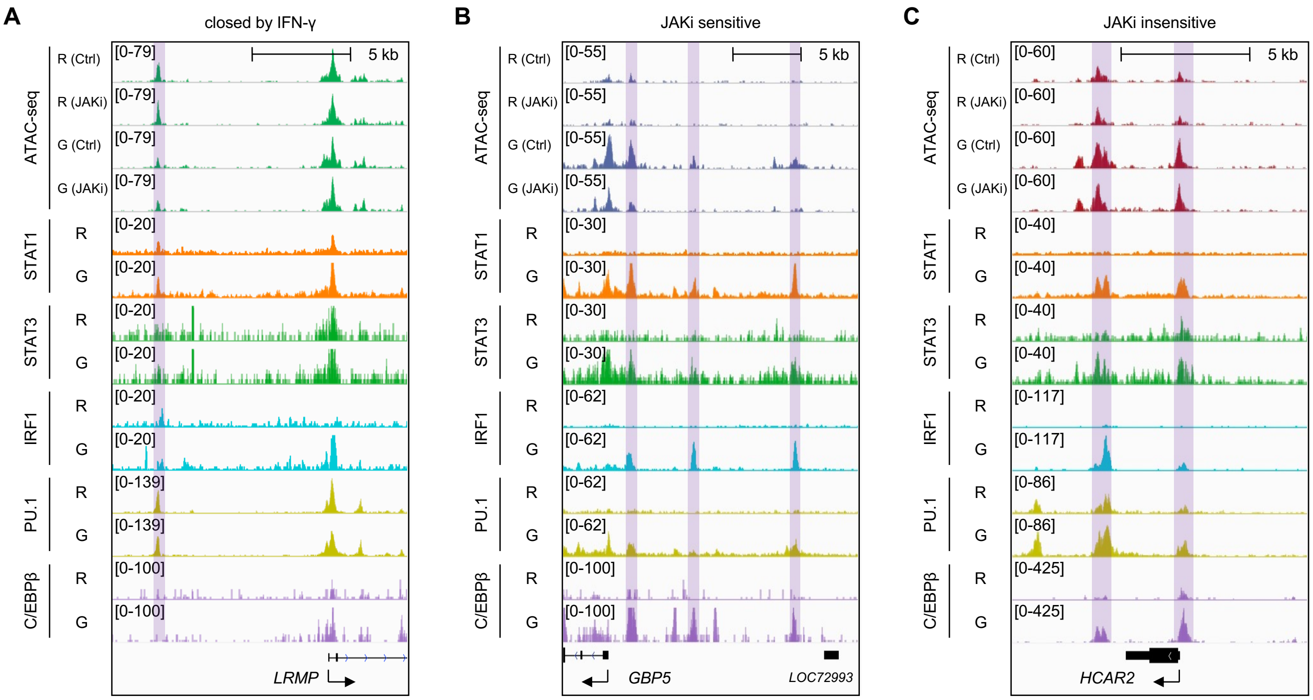

**Fig. S6. Binding of identified TFs to open chromatin regions regulated by JAKi.**

**(A-C)** The IGV Genome Browser displays ChIP-seq signals in the vicinities of indicated genes for chromatin remodeling regions. Clusters are indicated at the top. Clusters are indicated at the top. ChIP-seq and macrophage conditions are shown on the left. R, Resting macrophages; G, IFN-γ-primed macrophages.

**Supplementary Figure 7**

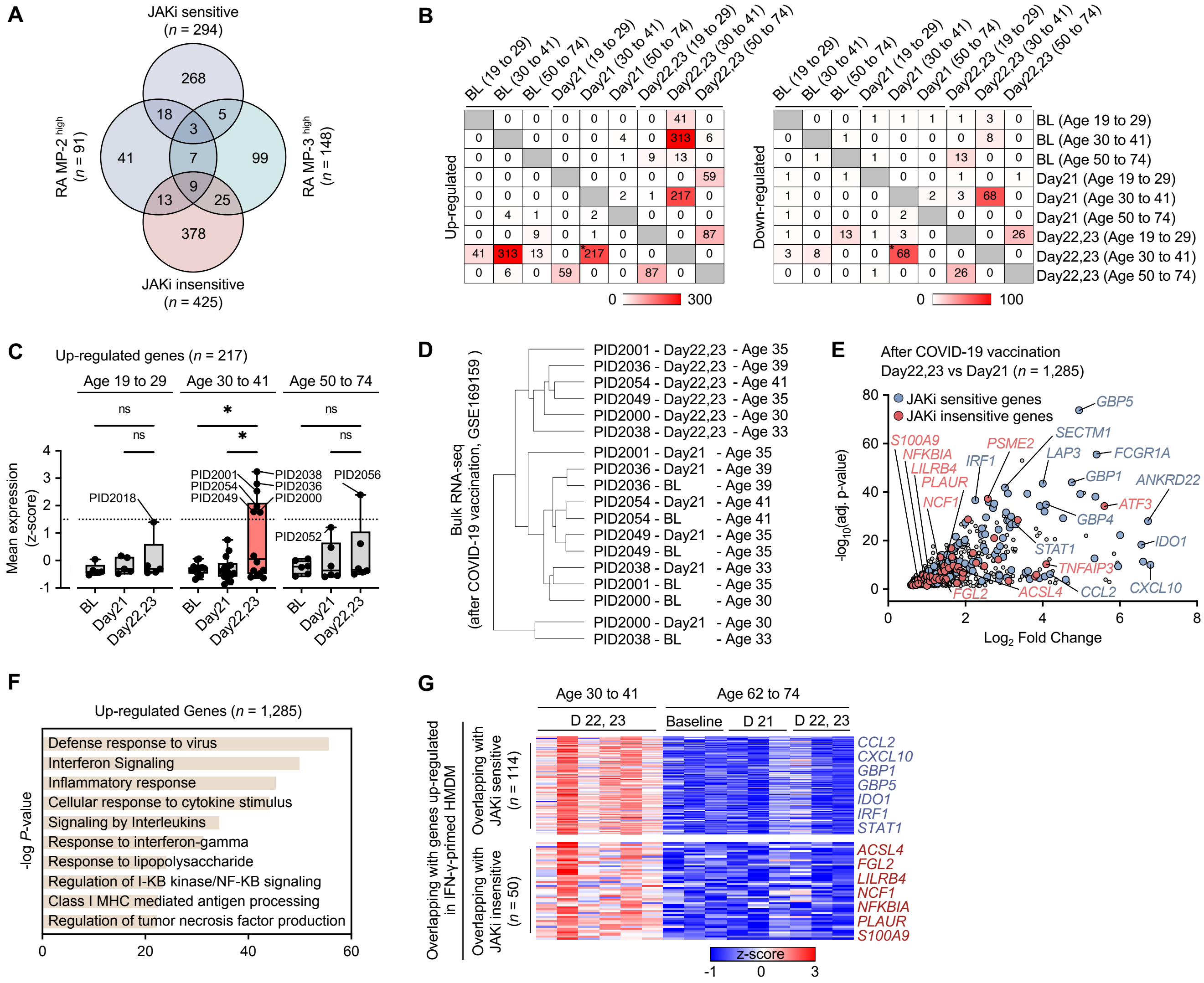

**Fig. S7. Identification of the JAKi-insensitive signature gene in inflammatory diseases.**

**(A)** Venn diagram to identify genes associated with IFN-γ-primed macrophages in RA MP-2 and RA MP-3 clusters. The JAKi-sensitive and JAKi-insensitive genes identified in IFN-γ-primed macrophages were used. **(B)** Analysis of all pairwise pairs of genes differentially regulated by the age group after COVID-19 vaccination. Heatmaps for the number of genes that are expressed differently by age. DE genes identified by EdgeR (FDR adjusted $P < 0.05$, fold change > 2) were used. TPM values of RNA-seq data were cut off to be greater than 4. Clusters are indicated on the left. RNA-seq data were obtained from GSE169159 (*, the combination with the highest total number of DE genes). **(C)** Bar plot of z-score mean values per patient for up-regulated genes (*n* = 217) at '22, 23 (age 30 to 41) versus 21 days (age 30 to 41)'. **(D)** Hierarchical analysis of gene expression for 6 selected patients. RNA-seq samples (*n* = 18) for days 21, 22 and 23 post-vaccinations, including baseline (BL), were used. **(E)** Scatterplot of genes upregulated in patients vaccinated against COVID-19. Colored dots indicate JAKi-sensitive or JAKi-insensitive genes. **(F)** Gene ontology (GO) analysis using genes up-regulated by COVID-19 vaccination. Bar plots showing the *P*-value (-Log10) significance of GO term enrichment for genes. GO analysis was performed using Metascape (http://metascape.org/). Up-regulated Genes, *n* = 1,285. **(G)** Age 62 to 74 heatmaps for JAKi-sensitive and JAKi-insensitive genes identified in the age 30 to 41 post-vaccination group. Clusters are shown on the left. The heatmaps for the age 62 to 74 group share the z-score values used for the age 30 to 41 group analysis Fig. 7D.

**Tables S1. Illumina/Nextera i5 common adapter and i7 index adapters.**

| Adapter name | Sequence (5' - 3') |
|---|---|
| Ad1_noMX | AAT GAT ACG GCG ACC ACC GAG ATC TAC ACT CGT CGG CAG CGT CAG ATG TG |
| Ad2.1_TAAGGCGA | CAA GCA GAA GAC GGC ATA CGA GAT TCG CCT TAG TCT CGT GGG CTC GGA GAT GT |
| Ad2.2_CGTACTAG | CAA GCA GAA GAC GGC ATA CGA GAT CTA GTA CGG TCT CGT GGG CTC GGA GAT GT |
| Ad2.3_AGGCAGAA | CAA GCA GAA GAC GGC ATA CGA GAT TTC TGC CTG TCT CGT GGG CTC GGA GAT GT |
| Ad2.4_TCCTGAGC | CAA GCA GAA GAC GGC ATA CGA GAT GCT CAG GAG TCT CGT GGG CTC GGA GAT GT |
| Ad2.5_GGACTCCT | CAA GCA GAA GAC GGC ATA CGA GAT AGG AGT CCG TCT CGT GGG CTC GGA GAT GT |
| Ad2.6_TAGGCATG | CAA GCA GAA GAC GGC ATA CGA GAT CAT GCC TAG TCT CGT GGG CTC GGA GAT GT |
| Ad2.7_CTCTCTAC | CAA GCA GAA GAC GGC ATA CGA GAT GTA GAG AGG TCT CGT GGG CTC GGA GAT GT |
| Ad2.8_CAGAGAGG | CAA GCA GAA GAC GGC ATA CGA GAT CCT CTC TGG TCT CGT GGG CTC GGA GAT GT |

**Tables S2. Primers for RT-qPCR.**

| Target gene | Primer | Sequence (5’ - 3’) |
|---|---|---|
| *TBP* | Forward | CCC GAA ACG CCG AAT ATA ATC C |
| | Reverse | AAT CAG TGC CGT GGT TCG TG |
| *CCL2* | Forward | AAG CAG AAG TGG GTT CAG GA |
| | Reverse | GCT GCA GAT TCT TGG GTT GT |
| *CXCL10* | Forward | GAA TCG AAG GCC ATC AAG AA |
| | Reverse | GCT CCC CTC TGG TTT TAA GG |
| *S100A9* | Forward | CAA AAT GTC GCA GCT GGA AC |
| | Reverse | CAC CAG CTC TTT GAA TTC CCC |
| *NFKBIA* | Forward | ACC TGG TGT CAC TCC TGT TGA |
| | Reverse | CTG CTG CTG TAT CCG GGT G |
| *PLAUR* | Forward | GAG CTA TCG GAC TGG CTT GAA |
| | Reverse | CGG CTT CGG GAA TAG GTG AC |
